# Spatial Flux Balance Analysis reveals region-specific cancer metabolic rewiring and metastatic mimicking

**DOI:** 10.1101/2024.11.28.625842

**Authors:** Davide Maspero, Giovanni Marteletto, Francesco Lapi, Bruno G. Galuzzi, Irene Ruano, Ben Vandenbosch, Ke Yin, Sabine Tejpar, Alex Graudenzi, Holger Heyn, Anna Pascual-Reguant, Chiara Damiani

## Abstract

To fully understand how cancer metabolism differs between primary tumors and metastases, resolving cell metabolism with spatial precision is essential. Yet, spatial fluxomics lags behind advancements in spatial transcriptomics.

To address this gap, we generated high-resolution spatial transcriptomics datasets from paired primary colorectal tumors and liver metastases, designed to capture metabolic adaptations across distinct tumor sites. Concurrently, we developed the Spatial Flux Balance Analysis (spFBA) computational framework to leverage them.

Since broad metabolic differences between tumors and healthy tissues are established, we first validated spFBA on a publicly available renal cancer dataset, including tumor-normal interface samples. spFBA detected cancer metabolic hallmarks, like enhanced glucose uptake and metabolic growth, but with unprecedented resolution, revealing lactate production with sustained oxygen consumption at the tumor interface and with reduced respiration in the core.

Next, applying spFBA to our colorectal cancer dataset, we provided biological insights, confirming that metastases mimic the metabolic traits of their tissue of origin. Additionally, our approach uncovered the first in vivo evidence of lactate-consuming cancer cells, marking a significant advance in understanding cancer metabolism.

spFBA stands out as a powerful approach to unravel the spatial metabolic complexity of cancer and beyond, leveraging the expanding landscape of spatial transcriptomics datasets.

## 2 Introduction

Cancer cells exhibit profound metabolic reprogramming to support their growth and survival, but these alterations are highly heterogeneous within and between tumors [1]. Differential wiring of the TCA cycle has been reported between naive and primed human stem cells [2], highlighting distinct metabolic activity also across different physiological cell states [2].

Although single-cell technologies have provided insight into the metabolic variability between cell-type clonal populations [3], they often overlook the critical role of spatial context. *In vivo* metabolism is influenced not only by intrinsic cellular factors but also by extrinsic cues such as nutrient gradients and interactions with neighboring cells. For example, stromal cells can interact with cancer cells and contribute to the local availability of metabolites [4].

To identify and compare spatially related metabolic phenomena across different tumor types, as well as between primary tumors and their metastases, an estimation of the metabolic fluxes with spatial resolution is essential to achieve this goal. The most direct approach to infer metabolic fluxes is Metabolic Flux Analysis (MFA) [5, 6], which involves iterative fitting of metabolite labeling patterns in isotopic tracers experiments. While spatial metabolomics techniques are developing rapidly, the progress of MFA lags far behind transcriptomics, at the bulk and especially at the single-cell, sub-cellular, and spatial levels [7, 8, 9]. This gap underscores the need to derive metabolic fluxes from gene expression data computationally. Although various factors, in addition to transcriptional regulation, contribute to fluxes and make this task inherently challenging, successful attempts have been made using bulk [10], single-cell [11, 12], or groups of biological samples, such as clusters of single cells identified through cluster analysis [13]. To our knowledge, no attempt has yet been made to model spatial spots or spatial clusters.

Building on prior work designed for bulk [14] and single-cell RNA-seq data [12, 15], we introduce spatial Flux Balance Analysis (spFBA). The spFBA framework processes spatial transcriptomics (ST) data to enrich metabolic fluxes at the spatial spot resolution. spFBA is rooted in the constraint-based (CB) modeling framework, the most established approach for flux simulation [16]. Although approaches based on self-supervised learning, which relax mass balance constraints, are emerging as a promising alternative to CB modeling [17], they do not provide information on reactions not directly linked to genes, such as oxygen uptake and the biomass synthesis pseudo-reaction, both of which are critical readouts in metabolic analysis.

To design spFBA, we selected the pipe-capacity approach [18], which retains the full network topology while adjusting flux boundaries based on expression data. This approach avoids the challenges of hard-threshold methods [19, 20], which risk overlooking nuanced variations in metabolic programs. Moreover, it does not assume that highly expressed reactions inherently carry more flux, as is assumed in expression-like flux optimization methods [21, 11]. A key feature of spFBA is its focus on relative expression differences across spatial spots rather than within individual spots. This design is grounded in the assumption that more abundant enzymes can carry greater flux, which is more reasonable when comparing the same enzyme across different spots, where catalytic efficiency is likely conserved.

Currently, the lack of spatially resolved flux measurements prevents the quantitative validation of spFBA predictions against ground-truth data. This limitation is a universal challenge in the field of metabolic flux inference, where validation efforts typically focus on specific, well-characterized pathways, such as glycolysis, rather than providing a comprehensive assessment of flux predictions. However, spatial datasets offer new opportunities to assess the global accuracy of flux predictions, as spatial patterns in metabolic fluxes are expected to emerge consistently with histological annotations. We devised a strategy to globally validate our results, which could serve as a validation framework for benchmarking future computational techniques. To let spatial patterns emerge from the data, we intentionally did not incorporate spatial coordinates into the spFBA problem, minimizing assumptions and parameters.

As a minimal preliminary check, we examined whether regions with similar gene expression patterns exhibit comparable flux distributions. It is reasonable to expect that cells with similar metabolic profiles activate corresponding metabolic genes. Although this outcome may seem intuitive, it cannot be assumed without verification, as steady-state constraints inherently reduce the degrees of freedom of flux distributions relative to gene expression data, unless a correlation between transcripts and flux is explicitly enforced as an objective function (e.g., as in [17]). Most importantly, recognizing that cancer tissues typically exhibit higher proliferation rates than adjacent normal tissues, we assessed whether the metabolic growth rates sustaining proliferation were indeed elevated in cancerous regions.

To test and apply spFBA, we focused on two tumor types originating from tissues that we hypothesize differ in nutrient availability: renal cell carcinoma (RCC) and colorectal cancer (CRC). Specifically, the microbial environment in the colon is likely to increase the availability of fermentation byproducts [22, 23]

For RCC, we selected a publicly available dataset that includes tumor-tissue interface samples from five different patients, ideal for our validation strategy. For CRC, we generated stereo-seq spatial datasets, having the high-resolution required to investigate tumor-stroma interactions *in vivo*. Importantly, this CRC dataset includes paired samples from primary and metastatic sites, suitable to investigate *in vivo* the recent hypothesis that the tissue of origin shapes the metabolism of metastases [24].

We demonstrate the capability of spFBA to delineate sustained metabolic growth in the tumor core compared to adjacent normal renal parenchyma. Using spFBA, we identified spatially distinct metabolic subpopulations within tumors, characterized by different nutrient consumption and secretion patterns. In RCC, spFBA recapitulated tissue architecture and captured hallmark features such as the Warburg effect, with pervasive lactate excretion in tumor cores. In CRC, spFBA revealed extensive lactate consumption by tumor cells in both primary and metastatic sites, alongside lactate production in stromal regions, consistent with the reverse Warburg effect.

## 3 Results

### 3.1 Spatial Flux Balance Analysis

We developed the spFBA framework, which takes as input any metabolic network reconstruction model and a sequencing-based spatial transcriptomics (ST) data to produce Flux Enrichment Scores (FES) for every reaction in each spatial spot.

spFBA is based on steady-state modeling and integrates the transcriptional information into flux boundary constraints. This approach enables the simulation of fluxes while accounting for indirect transcriptional regulation via substrate availability. spFBA enhances the expressive power of traditional pathway enrichment frameworks used in differential gene expression analysis by capturing differences in reaction directionality and metabolic growth rates, as well as in the preference for consumption or production of exogenous metabolites. As a result, a detailed and robust enrichment profile of fluxes across any reaction in the metabolic network used as input becomes available for each spatial spot within the ST dataset.

A key feature of spFBA is the use of spot-relative flux boundaries, which are defined by differential expression across spatial spots. This approach assumes that differences in enzyme abundance across spots correlate with flux capacity while maintaining catalytic efficiency as a constant. As a result, spot-relative models are meaningful only in comparative analyses, highlighting the interdependence of metabolic programs across spatial regions.

spFBA’s ability to generate a flux enrichment score for each spot independently, without explicitly using spatial coordinates, makes it possible to verify that transcriptionally similar spots also exhibit similar flux distributions Starting with a generic metabolic network model and information about nutrient availability, spFBA first uses Flux Variability Analysis (FVA) to define the region encompassing all flux distributions consistent with the steady-state assumption. Spatial transcriptomics data is then used to compute a Reaction Activity Score (RAS) for each reaction in each spot. Differences in the RASs across spots are used to determine subregions of the feasible flux space for each spatial spot (spot-relative model).

Once a spot-relative feasible region is obtained, in principle, it might be explored with any COBRA techniques, such as standard biomass optimization and parsimonious Flux Balance Analysis (pFBA). However, to release the optimality assumption, spFBA relies on flux sampling of the entire feasible region.

Finally, spFBA aggregates the sampled flux distributions into normalized centroids, delivering a final Flux Enrichment Score (FES) for each reaction in each spatial spot.

The overall spFBA framework is schematized in Figure 1.

**Figure 1.**
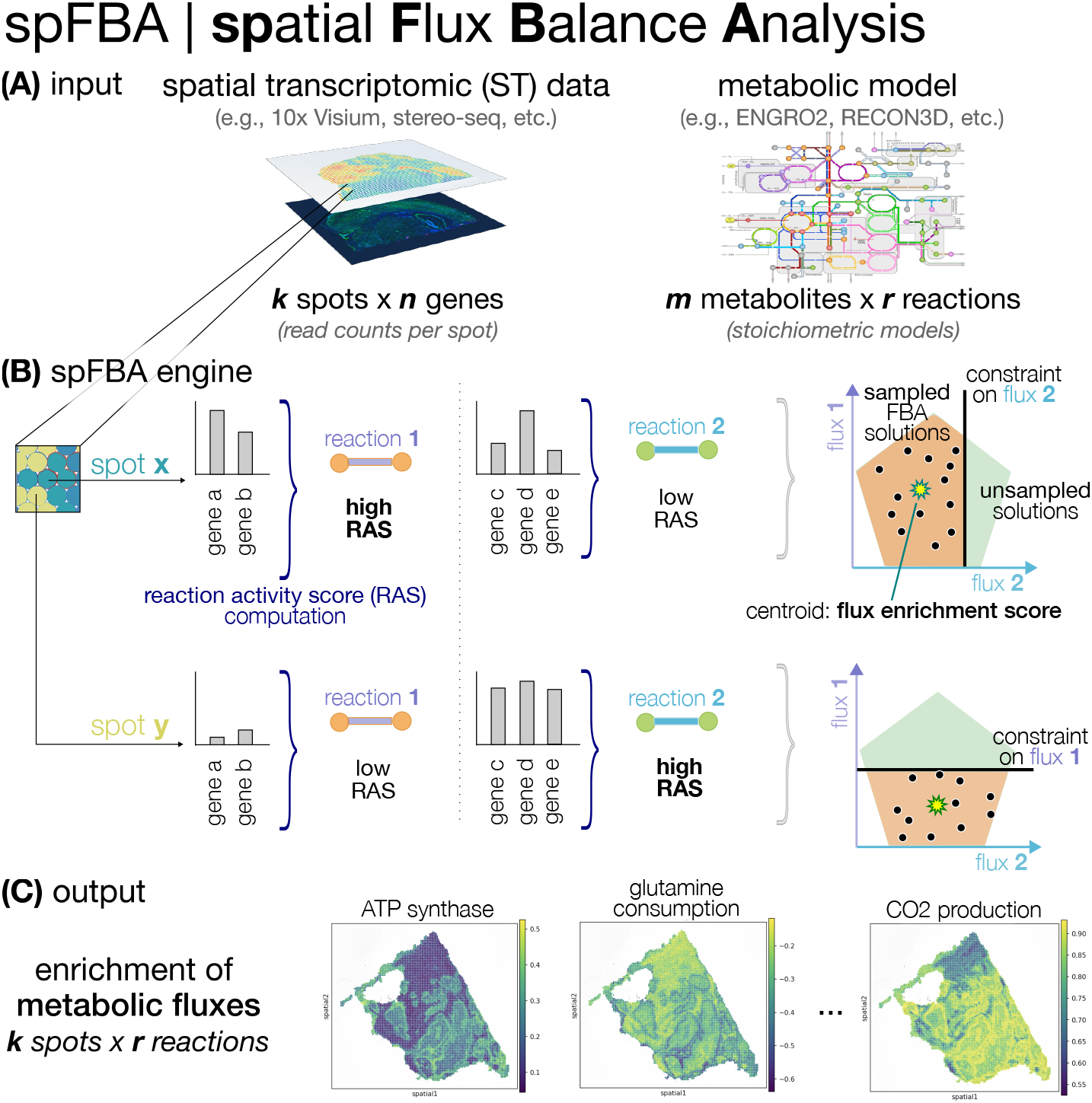
Overview of spFBA. **(A)** spFBA takes two data types as input: 1) a spatial transcriptomic (ST) read counts matrix of a biological sample, in the form of a *k* spots per *n* genes matrix; 2) a metabolic network reconstruction, either core (e.g., ENGRO2 [14]) or genome-wide (e.g., RECON3D [25]), in the form of a *m* metabolites per *r* reaction matrix. **(B)** After preprocessing via standard pipelines (see Methods), our framework computes for each reaction in each spot a reaction activity score (RAS), based on the expression level of the genes involved in the reaction. The RAS is then employed to set reaction-specific constraints in the Flux Balance Analysis (FBA) computation. Specifically, a high number of FBA solutions are randomly sampled in the constrained region and, for each reaction, the flux enrichment score is defined as the centroid of the distribution. **(C)** spFBA returns as output the flux enrichment scores, in the form of a *k* spots per *r* reactions matrix. Example heatmaps of three reactions are displayed, namely ATP synthase, glutamine consumption, and CO2 production.

From a computational perspective, spFBA is fully compatible with genome-scale metabolic models (e.g., RECON3D [25]). However, in this study, we opted to focus on central pathway models and employed the manually curated core model ENGRO2 [14] for its streamlined design and enhanced focus on key metabolic processes.

### 3.2 spatial FBA well recapitulates the tissue architecture

As an initial validation, we assessed whether the Flux Enrichment Scores (FESs) generated by spFBA exhibit a spatial distribution consistent with underlying histological structures. To this end, we compared clustering results obtained using three distinct feature sets representing different biological layers: gene expression, Reaction Activity Scores (RASs), and fluxes. To ensure fair comparisons, clustering parameters were independently optimized for each layer. The resulting clusters are visualized in Fig. 2 for interface samples and in Fig. S1 for core samples.

**Figure 2.**
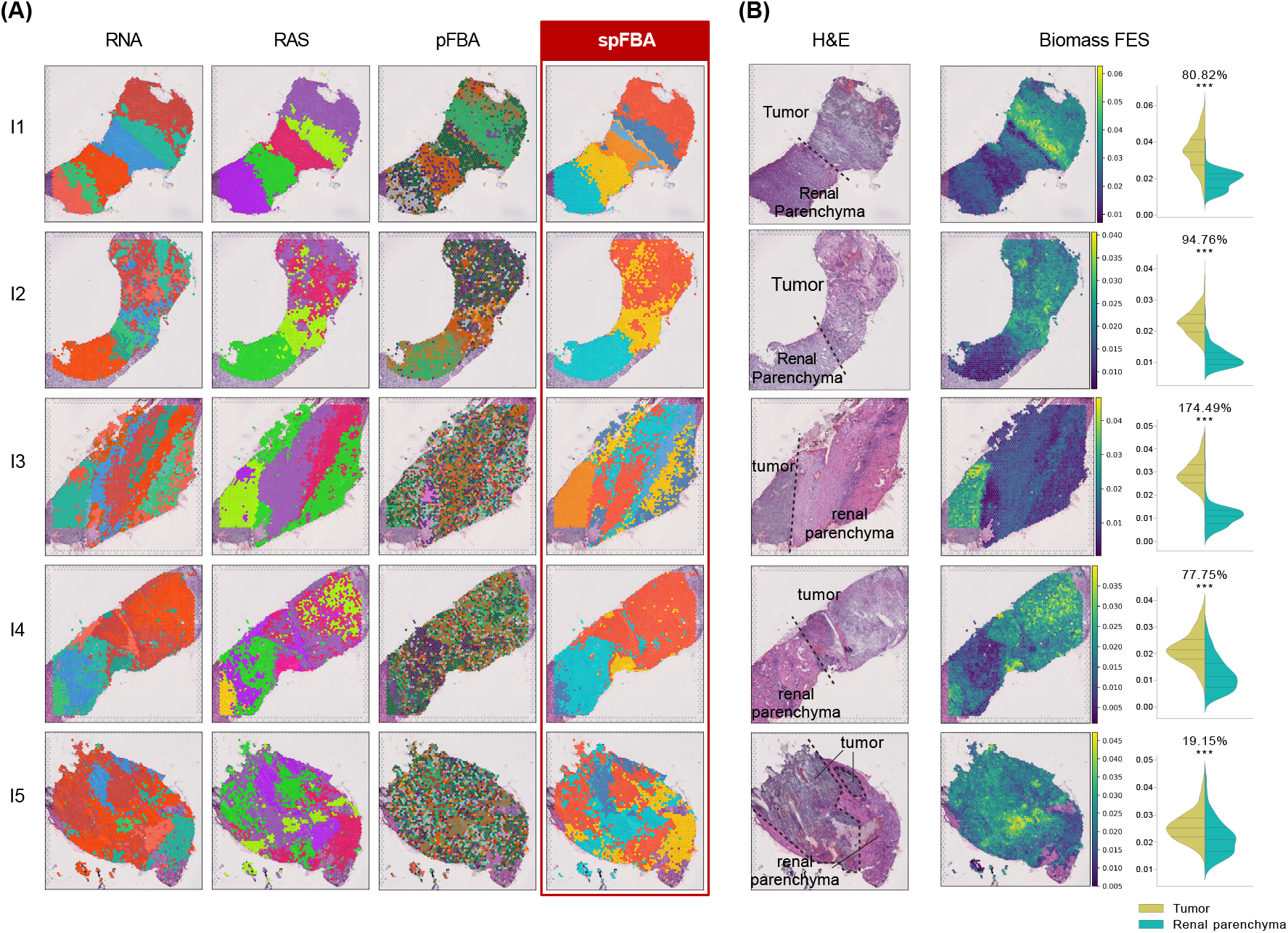
spFBA validation on ccRCC samples. **(A)** clustering results obtained with different data layers: RNA (preprocessed reads counts), RAS, pFBA and spFBA. Color scales were independently assigned to clusters, so they are not comparable across layers or samples. **(B)** From left to right: annotated H&E staining taken from [26]; Biomass FESs for each spot obtained with spFBA; Statistical comparison of the biomass FESs between spots annotated as tumor and parenchyma. The violin plot titles indicate the percentage of variation in the averages of the two populations and the results of t-tests. Statistical significance is reported using a star notation system, with a significance threshold of 0.05. When applicable, the same information for the ccRCC tumor core samples are in supplementary Figures S1.

Our analysis demonstrated that clusters emerging from the FESs exhibit a coherent spatial distribution that closely aligns with the histological architecture revealed by the transcriptomics layer. Importantly, these FES solutions were generated independently for each spatial spot, relying solely on transcriptomic content and without explicit consideration of spatial coordinates. It can however be noticed that the number of clusters observed in the fluxomics layer is generally lower than that in the transcriptomics layer. This outcome is expected due to the reduced degrees of freedom inherent to the steady-state fluxome compared to the transcriptome, as previously reported in [12].

To demonstrate that the observed alignment is not a trivial outcome, merely resulting from trascriptionally-dependent flux boundaries, we repeated the analysis using the same spot-relative models, thus incorporating the same trascriptionally-dependent boundaries, but optimizing them for biomass production. As shown in Fig. 2A, when fluxes obtained with parsimonious Flux Balance Analysis (pFBA) were used as clustering features, instead of our the flux-sampling derived FESs, the resulting clusters were way more scattered and less reflective of the underlying histological organization. A similar pattern was observed in tumor core samples (Fig. S1). To further support these findings, we performed a quantitative comparison of clustering outcomes across layers for all clear cell Renal Cell Carcinoma (ccRCC) samples. Using the average V-measure as a metric, we observed that normalized sampling centroids consistently preserved spatial coherence more effectively than pFBA-derived fluxes (see Table S1).

This analysis highlights that, while FES-based clustering aligns well with histological structures, this alignment is not inherent and depends on the approach used.

### 3.3 spatial FBA detects cancer enhanced metabolic growth

To verify that metabolic growth rates are enriched in the tumor region, we used the Flux Enrichment Score of the biomass pseudo-reaction as a proxy. We compared the biomass FES between tumor regions and renal parenchyma in interface samples from five patients profiled in [27]. Hematoxylin and Eosin (H&E) staining images provided by the authors were annotated to delineate tumor and healthy tissue regions for samples I1 and I2, where the tumor-normal boundary is most distinct. While the authors had annotated tumor and healthy tissue regions for samples sI1 and I2, where the tumor-normal boundary is most distinct, we extended these annotations to the remaining interface samples. Fine-grained pathological annotations to accurately exclude non-tumoral regions were not feasible. This limitation reflects the constraints of working with low-resolution datasets, where distinguishing fine structural details is challenging.

Figure 2B presents the annotated H&E staining alongside the biomass FESs for the interface samples and the frequency distributions of spots annotated as tumor or normal. The data reveal a clear and statistically significant separation between the two regions, with higher biomass enrichment consistently observed in the tumor region.

To further demonstrate the added value of simulation-based enrichment, we qualitatively compared the biomass FES with cell-cycle scores, which estimate the most likely phase representative of each spot based on the expression of predictor genes for cell-cycle phases [28]. As shown in Supplementary Fig. S2, the biomass FES calculated by spFBA provides superior distinction between tumor cores and renal parenchyma compared to the cell-cycle score, underscoring the advantage of spFBA over transcriptomic approaches in characterizing metabolic behavior.

### 3.4 spFBA captures the Warburg effect in ccRCC

Using spFBA, we characterized spatially resolved metabolic reprogramming in ccRCC, focusing on the Warburg effect. This metabolic phenotype, described a century ago [29], involves preferential energy generation through glycolysis followed by lactate fermentation, even in the presence of oxygen.

In addition to glucose metabolism, we examined the spatial distribution of other key metabolites, including glutamine [30], glutamate [30], serine [31], glycine [32], and palmitate, which has been implicated in metastasis [33]. Fig. 3A shows the spatial distribution of the FESs for the reactions under study in a representative interface sample (I2). Panel B of the same figure presents the statistical comparison of these FESs between spots annotated as tumor and renal parenchyma, across all interface samples.

**Figure 3.**
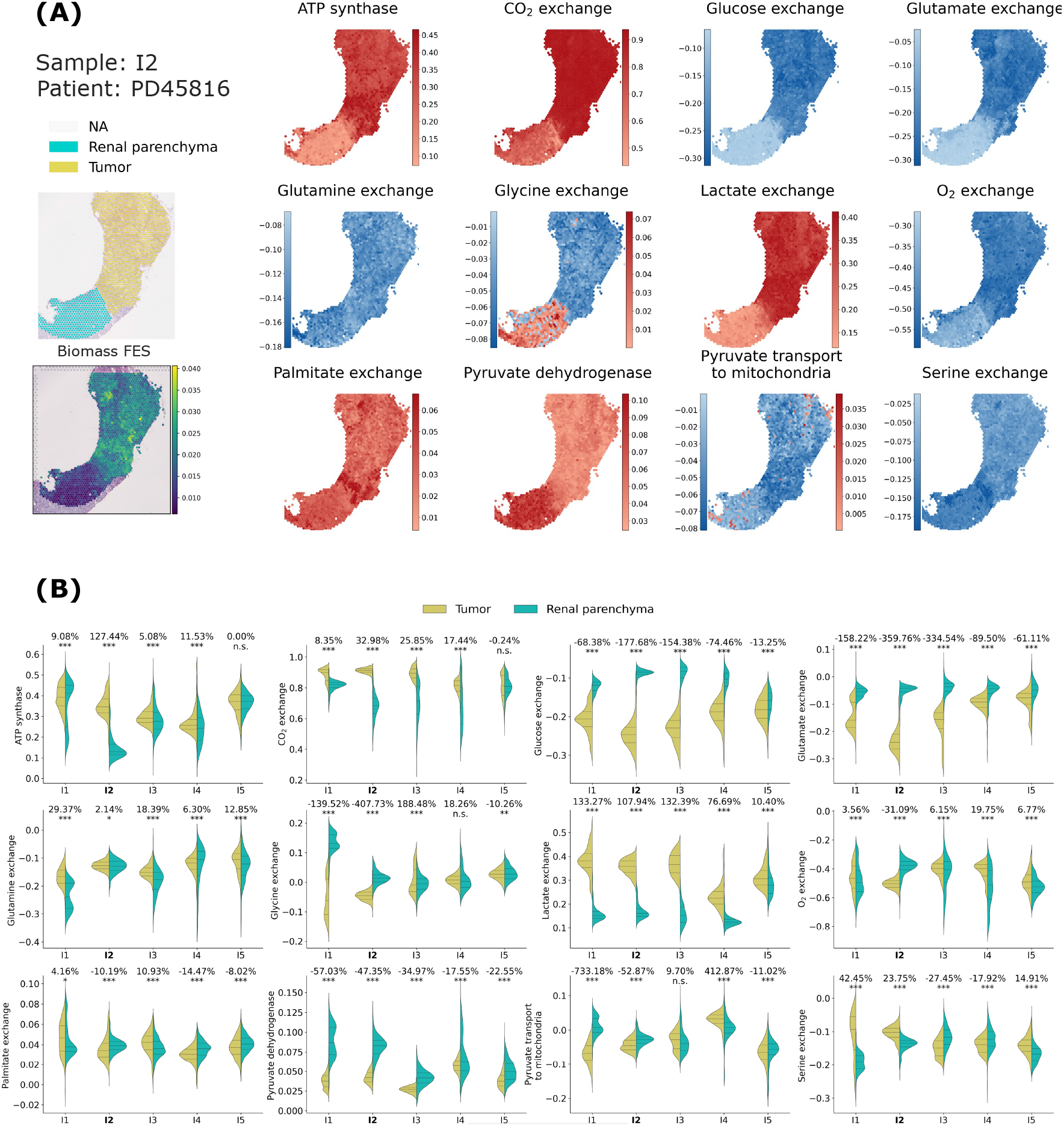
Tumor vs. parenchyma **(A)** The FES of a set of reactions of interest (in alphabetic order) is visualized for ccRCC sample I2. In the case of exchange reactions, negative values correspond to consumption of the metabolite, positive values to production. **(B)** Statistical comparison of the FESs for the reaction in panel A between spots annotated as tumor and parenchyma for all ccRCC interface samples.The violin plot titles indicate the percentage of variation in the averages of the two populations and the results of t-tests. Statistical significance is reported using a star notation system, with a significance threshold of 0.05.

A striking observation is the markedly higher glucose consumption and lactate production in the tumor region (corresponding to the upper area, as per annotation in the top left corner in Fig. 3A), as expected under the Warburg effect. This difference in glucose consumption and lactate production rates was substantial and statistically significant consistently in all interface samples (Fig. 3B). The specific spatial distribution of the other samples is reported in Fig. 4 for patient PD47171 samples and in the supplementary material for all the other kidney samples (Figs. S3 to S9).

**Figure 4.**
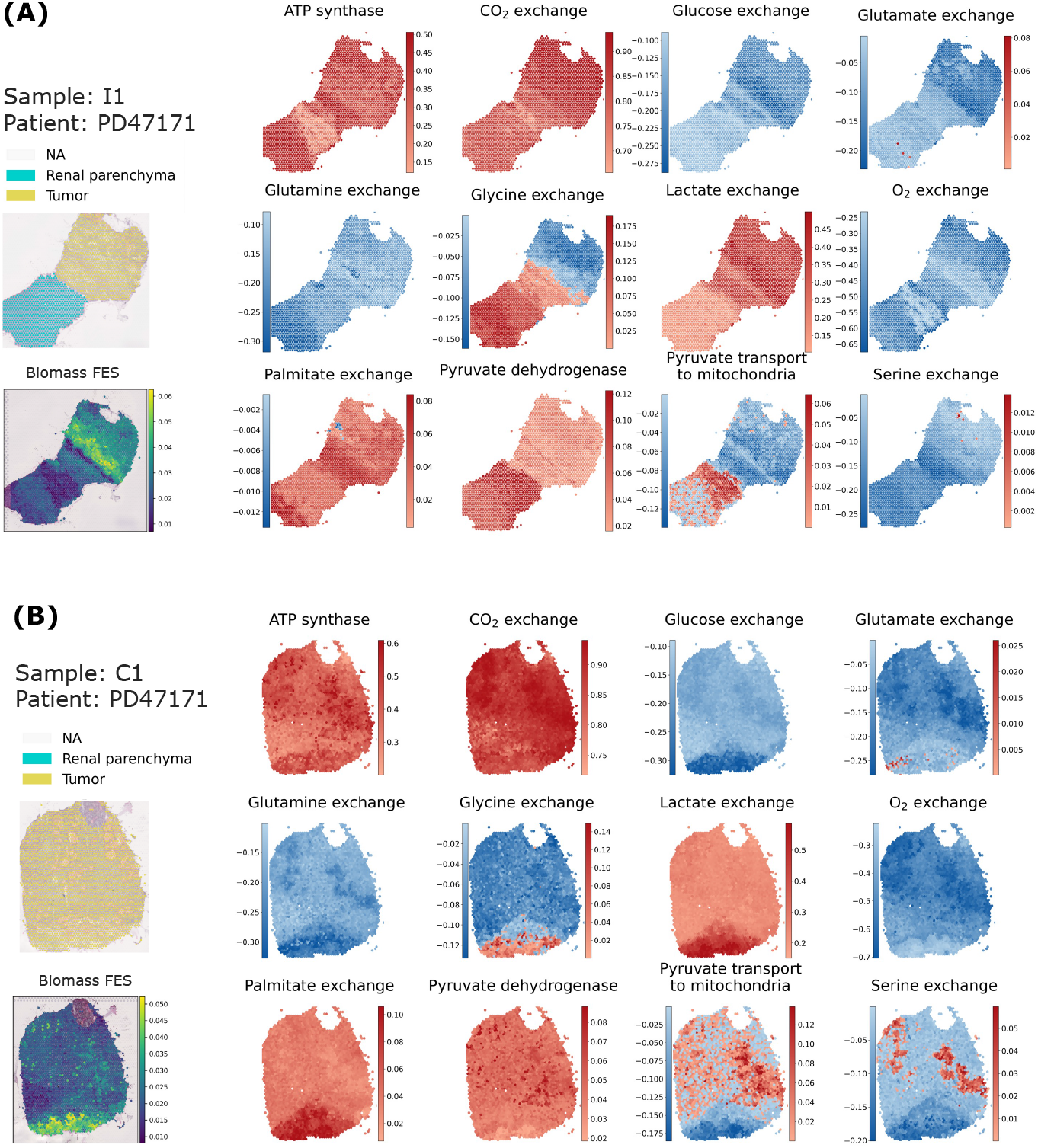
Paired interface and tumor core samples. The FES of a set of reactions of interest (in alphabetic order) is visualized for ccRCC interface sample I1 (panel A) and tumor core sample C1 (panel B). In the case of exchange reactions, negative values correspond to consumption of the metabolite, positive values to production.

We remark that, conventionally, intake fluxes are represented with a negative sign, meaning that a more positive distribution reflects lower consumption. Interestingly, oxygen consumption rates are generally comparable between the tumor and normal regions. In most samples, oxygen consumption appears just slightly reduced in tumor regions, with decreases ranging from 3% to 20%. However, the distributions are far from normal, and these differences do not seem statistically robust. In contrast, in sample I2, oxygen consumption shows a notable increase of approximately 30% in the tumor region. This pattern of oxygen consumption, alongside enrichment in other metabolic indicators —such as *CO*_2_ production and ATP synthesis —suggests that cancer cells adopt a hybrid respiratory-fermentative metabolism. This supports the idea that cancer cells are generally more metabolically active and ‘greedy’ (Fig. 3). One might be tempted to interpret lactate production in this scenario as a simple result of oxygen limitation, where excess glucose that cannot be oxidized is fermented into lactate. However, the situation portrayed by spFBA is far more complex. To explore the interplay between glycolysis and mitochondrial activity, we examined the FES for pyruvate dehydrogenase, which controls the entry of pyruvate into the TCA cycle. Remarkably, pyruvate dehydrogenase activity is significantly reduced in tumor regions across all interface samples, indicating that glucose-derived pyruvate is not oxidized via the TCA cycle. Instead, pyruvate transport to the mitochondria is negligible in the renal parenchyma and becomes negative in tumor regions, suggesting a reversal of transport from mitochondria to the cytosol. However, the frequency distribution of the FES for this reaction straddles zero, indicating that negative values may result from probabilistic variation rather than definitive biological reversal. In sample I2, the analysis of the confidence interval for the FESs reveals that the lower and upper bounds (Fig. S10 and S11) are nearly identical across most regions, indicating minimal uncertainty in the estimates. Nevertheless, a few scattered spots in the normal region show subtle shifts in the pyruvate transport FES from blue to red. This suggests that while reverse transport is generally negligible, localized variations may occasionally result in fluxes being classified as negative instead of positive.

Although this effect is limited and does not change the broader conclusion that cancer cells bypass canonical mitochondrial pathways, it underscores the need to carefully interpret fluxes near zero.

### 3.5 spFBA unveils distinct variants of the Warburg Effect across regions of the same tumor

The situation portrayed by scFBA at the tumor-normal interface aligns more closely with the “selfish cell” interpretation of the Warburg effect [34], where cancer cells prioritize nutrient consumption to fuel their metabolic needs without repressing the respiratory chain.

In [35] it was theoretically shown with a simple metabolic network model that, if available oxygen is not sufficient to oxidize available carbon sources fully, utilization of glutamine via reductive carboxylation and conversion of most glucose to lactate confer an advantage for biomass production. spFBA portrayed a similar situation of high oxygen consumption coupled with low pyruvate dehydrogenase activity emerging from the ST data, with the more detailed metabolic network ENGRO2, which includes all essential and non-essential amino acids. Interestingly, spFBA revealed that, across all interface samples, glutamine is not enriched in tumor regions one might expect. Instead, its downstream metabolite, glutamate, shows significant enrichment (Fig. 3).

The selfish cell interpretation of the Warburg effect contrasts with the Crabtree effect [36], which involves a suppression of oxidative phosphorylation. However, we identified metabolic patterns consistent with the Crabtree effect in tumor core samples. For example, when comparing the interface sample I1 (Fig. 4A) with its paired tumor core sample C1 (Fig. 4B), we noticed that the bottom region of the tumor core exhibits a more fermentative phenotype. This region shows elevated lactate production and higher glucose intake, but reduced oxygen consumption compared to the upper core region. Notably, this fermentative area also displays enhanced biomass production, indicative of a Warburg-like phenotype.

Interestingly, differently from the tumor interface (Fig. 4A) glutamate consumption is not enriched in this high-growth area. Instead, there is a tendency to secrete exogenous glutamate, while glutamine consumption becomes more pronounced. This metabolic shift suggests a localized adaptation to support growth under varying environmental conditions.

Collectively, these findings reveal a complex metabolic landscape in ccRCC, with distinct Warburg phenotypes coexisting in different tumor regions. At the interface, the Warburg effect appears to coexist with oxidative phosphorylation, consistent with the hybrid metabolic phenotype described in some cancer cells [37]. In contrast, the tumor core displays regions with more classical fermentative behaviors, akin to the Crabtree effect. These observations underscore the spatial and functional heterogeneity of cancer metabolism, highlighting the adaptive strategies tumors employ to sustain growth and proliferation.

### 3.6 scFBA can enrich metabolic interactions

A key advantage of scFBA over standard enrichment frameworks is its ability to identify spatial spots exhibiting opposing signs for exogenous nutrients — negative for consumption and positive for production. This feature is critical for uncovering metabolic cooperation within tissues.

For instance, in both interface sample I1 and its paired tumor core sample C1 (Fig. 4), as well as in interface I2, distinct regions can be observed where glycine is either produced or consumed. Specifically, one region tends to produce this metabolite, while another tends to consume them.

This phenomenon is particularly pronounced in the tumor core C1, where the glycine-producing region overlaps with an area of lower oxygen consumption. This overlap suggests a potential link between glycine production and metabolic adaptation to the tumor microenvironment, such as reduced oxygen availability.

While serine also shows regions with opposing metabolic signs in C1 (Fig. 4), the positive regions (highlighted in red) overlap with large congested blood vessels and multifocal areas of hemorrhage, as indicated by H&E staining (Supplementary Fig. S1). These histological features may affect the reliability of the results, potentially reflecting tissue-specific artifacts rather than genuine metabolic patterns.

It is important to note that we cannot confirm whether these populations exchange nutrients among themselves or only with the plasma. Determining this would require experimental measurements of plasma nutrient consumption and secretion rates, which are challenging to obtain *in vivo*. Additionally, a population mass balance constraint must be applied, as suggested in [12]. However, the absence of regions exhibiting lactate consumption enables us to rule out metabolic cooperation involving lactate in all ccRCC samples.

Notably, with only a few exceptions, most regions across all ccRCC samples do not consume exogenous lipids. Instead, they tend to secrete them, indicating a reliance on de novo lipogenesis — even in excess — to meet the metabolic demands of growth, as reported for many cancer cells [38].

### 3.7 CRC liver metastasis mimics tissue-of-origin metabolic traits

When applying spFBA to our CRC samples, we used the very same experimental settings as for ccRCC, ensuring that any differences observed in the results are solely due to variations in the spatial transcriptomics information. It is particularly striking that, while lactate secretion was consistently detected by spFBA in all ccRCC samples, none of the CRC samples showed any lactate secretion (Fig. 5, S14,S13). On the contrary, most regions in all CRC samples exhibited an enriched lactate uptake.

**Figure 5.**
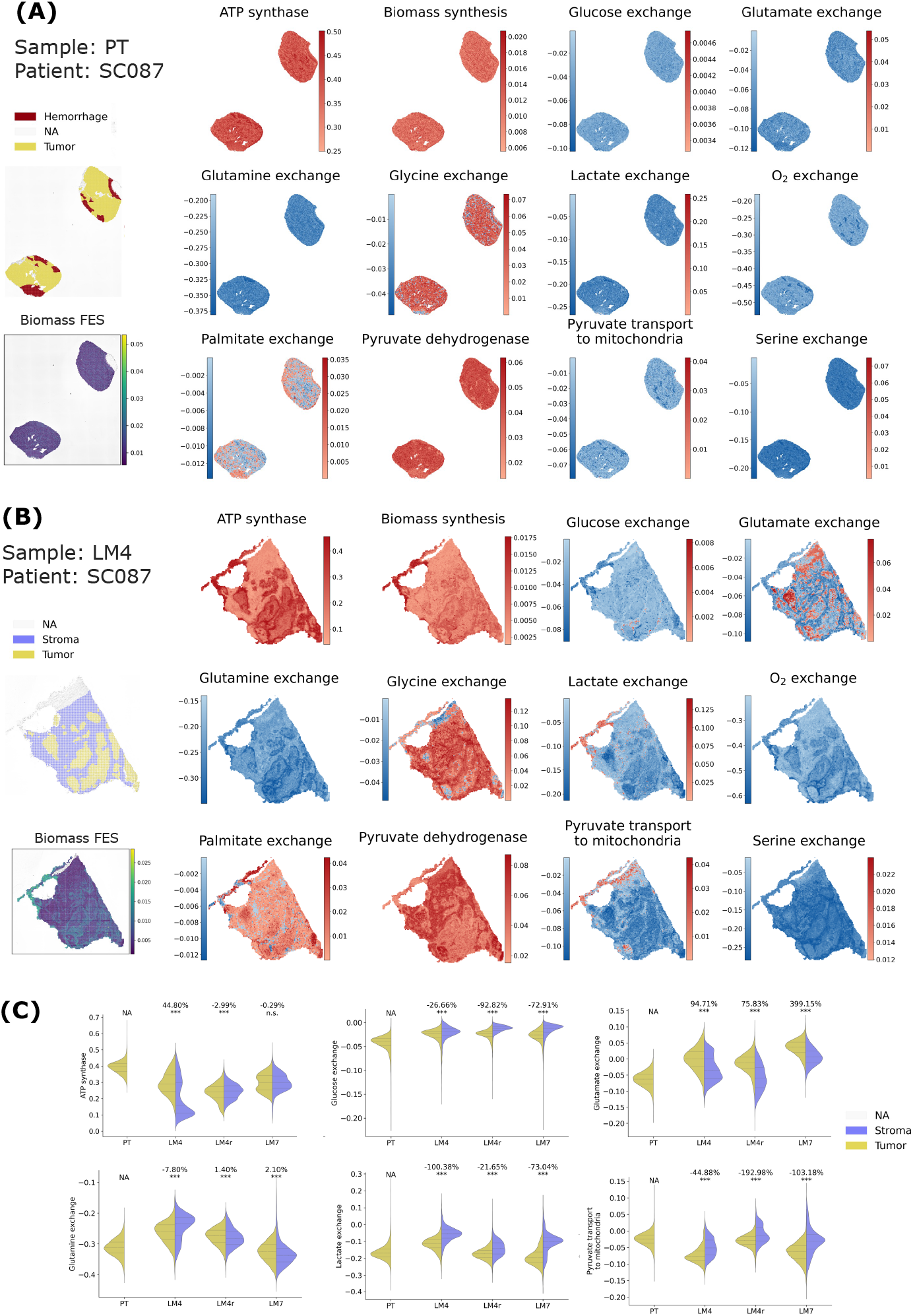
**(A) and (B)** Primary tumor vs. metastasis. FESs of a set of reactions of interest (in alphabetic order) for CRC PT (panel A) and paired LM4 sample (panel B). In the case of exchange reactions, negative values correspond to consumption of the metabolite, positive values to production. **(C)** Tumor versus stroma. Statistical comparison of the FESs for a subset of the reactions in panel B between spots annotated as tumor and stroma, for all CRC samples. When applicable, the violin plot titles indicate the percentage of variation in the averages of the two populations and the results of t-tests. Statistical significance is reported using a star notation system, with a significance threshold of 0.05. The same information for all reactions in panel B is in Fig. S12

We hypothesize that more exogenous lactate is available for both tumor and non-tumor cells to consume in the colon compared to the kidney. This may be due to the presence of various bacteria in the colon capable of producing lactate [39]. In contrast, in the kidney, exogenous lactate is likely less abundant, as supported by literature reporting that kidney cells tend to produce rather than consume lactate [40].

Remarkably, lactate uptake was observed both in CRC primary tumor (PT) and in liver metastasis (LM) (Figs. 5, S14,S13). The liver is one of the main organs in charge of lactate metabolism, which presents a net uptake of lactate [41]. Therefore, enhanced lactate consumption by metastatic tumor cells might be supported by the pre-existent context of lactate accumulation in physiological conditions. The common utilization of lactate by PT and LM highlighted by scFBA supports the recent hypothesis that differences in tissue nutrient availability might constrain the tissue-of-origin-shaped metabolism of cancer cells to limit sites of metastatic colonization [24].

Metastatic migration of colon cancer cells could depend on lactate accessibility of the destination tissue, which enhances the metabolic plasticity of tumor cells to react to changes in nutrient availability, thus maximizing cellular proliferation and growth.

### 3.8 The reverse Warburg effect takes place in CRC liver metastasis

High-resolution ST allowed us to identify differences in metabolic fluxes between tumor cells and associated stroma. The annotated CRC H&E stained samples are reported in Supplementary Fig. S15. Regarding lactate metabolism, the FES of lactate exchange for the LM in Fig. 5B highlights dark blue patches that correspond to tumor cells, and ligher areas surrounding them that represent the stromal tissue. This contrast indicates that tumor cells present enhanced lactate consumption, whereas some stromal cells seem to be producing lactate. The statistical comparison of the spots annotated as stroma and tumor reported in Fig. 5C confirms that lactate consumption is enriched in the tumor regions in all CRC samples. On the contrary, it is very poor in stromal regions, which occasionally secrete lactate. Additionally, those stromal regions also showed decreased glucose consumption, but increased glutamate consumption. Histologically, stromal regions surrounding the tumor are composed of abundant mature connective tissue with organized cancer-associated fibroblasts (CAFs). Tissue morphology together with spFBA findings support the “reverse Warburg effect” phenomenon, in which CAFs glycolysis metabolically supports adjacent cancer cells [4]. In conclusion, the preexistent context of lactate accumulation exploited by metastatic tumor cells in normal hepatic parenchyma might be further increased by lactate production by CAFs located in the stroma.

It is worth noticing that the reverse Warburg effect is more evident in liver metastases than in primary colon tumors. In the latter, the absence of clearly annotated stromal areas and the predominance of cancer cells result in uniformly high lactate consumption, without the distinct spatial heterogeneity observed in metastatic samples Importantly, spFBA is shedding new light on the reverse Warburg effect. In particular, it can be observed that the pyruvate that necessarily derives from lactate consumption is not used to fuel the TCA cycle. On the contrary, pyruvate transport is mainly enriched from mitochondria to the cytosol, indicating that lactate-derived pyruvate must be employed in biosynthetic pathways within the cytosol.

### 3.9 Spatial distribution of metabolite exchange highlights tumor-stroma interface activity

In the previous section, the violin plots comparing tumor and stromal regions revealed significant differences in some metabolic fluxes. However, for certain reactions—such as biomass production or oxygen consumption—these differences appeared less pronounced. This limited contrast is likely driven by intratumoral heterogeneity, rather than a uniform metabolic profile across the tumor region.

Spatial mapping through spFBA uncovers this heterogeneity, demonstrating that metabolic activity is not evenly distributed within tumor regions. As noticeable in Fig. 5 and better highlighted in Fig. S16, glutamate exchange exemplifies this phenomenon. While overall glutamate consumption is lower in tumor regions compared to the stroma, spatial visualization reveals distinct metabolic behaviors at the tumor-stroma interface. Black arrows indicate that glutamate production is concentrated at the tumor periphery, where cells are in direct contact with stromal regions. In contrast, cells within the tumor core predominantly consume glutamate, albeit at a lower rate than stromal cells. Notably, this pattern is absent in primary tumor regions that lack stromal interaction. In primary tumors, glutamate is consistently consumed across the entire region, and no peripheral production is observed. This suggests that glutamate production may be specifically induced by the tumor’s interaction with stromal cells, highlighting a possible metabolic adaptation at the invasive front. Worth of note, according to the literature, glutamate exchange could be related to acquired drug resistance or epithelial-mesenchymal transition [42]

These findings underscore the advantage of spatial data, as single-cell analyses alone would only capture aggregate differences in metabolite consumption or production. By visualizing where metabolite exchange occurs, spFBA coupled with high resolution provided by stereo-seq data provides a more nuanced understanding of intratumoral metabolic heterogeneity and identifies regions at the tumor-stroma interface as potential hotspots of metabolic rewiring.

## 4 Conclusions

We introduced Spatial Flux Balance Analysis (spFBA), a computational framework to characterize metabolism with spatial resolution by leveraging transcriptomics data.

spFBA demonstrated a unique capability to uncover the spatial metabolic heterogeneity underlying cancer evolution. It confirmed well-established cancer metabolic hallmarks, such as enhanced metabolic growth and the Warburg effect, achieving unprecedented resolution at the level of individual metabolic reactions. This resolution enabled the identification of spatially distinct tumor subpopulations with specific flux distributions.

While the Warburg effect is often discussed as a singular phenomenon, spFBA reveals spatial heterogeneity in its manifestation within the same tumor. At the tumor-normal interface, the Warburg effect aligns with the “selfish cell” model [34], where glucose is converted to lactate without suppressing respiratory chain activity. In contrast, tumor core samples display distinct metabolic profiles, including patterns resembling the Crabtree effect [36].

Unlike standard enrichment approaches that identify active pathways, spFBA provides a deeper insight into the directionality of metabolic fluxes, offering a more nuanced understanding of tumor metabolism. For instance, while enrichment analysis could identify lactate metabolism alterations, spFBA revealed tissue-specific metabolic strategies, such as lactate uptake in colorectal cancer (CRC) compared to secretion in renal cell carcinoma (RCC), indicating distinct metabolic adaptations.

When applied to a new high-resolution colorectal cancer dataset, including primary tumor and matched liver metastasis samples from the same patient, spFBA provided the first clear in vivo evidence of lactate-consuming subpopulations. These findings align with the reverse Warburg effect previously observed in certain cancer cell lines, now revealed with unprecedented spatial detail.

Beyond reproducing known phenomena, spFBA revealed novel metabolic behaviors, such as the decoupling of lactate uptake from oxidative fluxes, opening new avenues for experimental validation and therapeutic intervention. Furthermore, it demonstrated a remarkable similarity between the metabolic behavior of primary CRC tumors and their metastases, supporting the hypothesis that nutrient availability in the tissue of origin – such as the high levels of lactate in the gut – shapes the metabolic phenotype of metastatic cells.

It is important to acknowledge certain limitations of our approach. The framework relies on a probabilistic distribution to infer metabolic fluxes, providing a relative enrichment based on the expected value rather than an absolute measure of flux. This means that scFBA does not quantify the magnitude of flux but instead indicates whether it is more or less active compared to a reference cell population. Additionally, spFBA estimates metabolic activity based on probability distributions and expected values. Although confidence intervals can help address uncertainty around the expected value, no model can guarantee that cells operate at the expected (most probable) metabolic state, as less likely states may still occur. In this regard, spFBA shares the limitation inherent to all probabilistic approaches.

Despite these limitations, scFBA remains a powerful tool not only for studying tumor heterogeneity and progression but also for exploring metabolism across a wide range of tissues and diseases. Importantly, spFBA can be seamlessly applied to the growing wealth of sequencing-based spatial transcriptomics datasets, enabling the fine characterization of metabolic processes without the need for additional experimental data.

## 5 Methods

### 5.1 Experimental data

#### 5.1.1 ccRCC dataset

We analyzed a publicly available spatial transcriptomics dataset[43], where kidney samples from 7 patients with Clear Cell Renal Cell Carcinoma (ccRCC) were sequenced using the 10x Visium Genomics protocol.

The patients underwent surgical resection of radiologically diagnosed, treatment-naive renal tumors. Most of the patients were in significantly advanced stages of the disease, we refer the reader to [26] for further details. Among the 16 samples available, we kept 5 samples from the tumor core and 5 from the interface between primary tumor and healthy tissue with higher quality considering metadata annotations provided by the original authors. See table 1.

**Table 1:** Dataset metadata. Additional information regarding patient, sample and quality metrics are reported for each dataset.

| Sample name | Sample ID | Patient ID | Region | Spot area [$\mu m$] | Spot # | Unique genes | Mean counts per spot | Mean genes per spot |
| --- | --- | --- | --- | --- | --- | --- | --- | --- |
| LM4 | A02991A2 | SC087 | Liver met. | 625 | 9539 | 24673 | 3526.04+-1290.24 | 1630.53+-512.48 |
| LM7 | C03445C6 | SC087 | Liver met. | 625 | 20729 | 25425 | 2569.98+-812.26 | 1348.27+-362.47 |
| PT | C03445G5 | SC087 | Colon primary tum. | 625 | 14956 | 25688 | 5592.64+-2137.28 | 2224.24+-562.71 |
| LM4r | A03389C5 | SC087 | Liver met. | 625 | 11919 | 24974 | 4912.33+-1389.13 | 1941.12+-479.57 |
| C1 | 6800STDY12499410 | PD47171 | Kidney - Tum. Core | 2376 | 2888 | 22317 | 5821.02+-3532.71 | 2334.16+-1002.04 |
| C5 | 6800STDY12499413 | PD45816 | Kidney - Tum. Core | 2376 | 2102 | 19735 | 2776.64+-1891.10 | 1193.50+-549.47 |
| C2 | 6800STDY12499407 | PD43948 | Kidney - Tum. Core | 2376 | 832 | 19822 | 10290.89+-5756.34 | 3065.35+-1101.38 |
| C4 | 6800STDY12499412 | PD45814 | Kidney - Tum. Core | 2376 | 1026 | 21488 | 9650.70+-4667.24 | 3324.03+-1184.36 |
| C3 | 6800STDY12499408 | PD47512 | Kidney - Tum. Core | 2376 | 3800 | 22184 | 3323.62+-2214.55 | 1778.00+-779.10 |
| I4 | 6800STDY12499502 | PD45814 | Kidney - Tum. Interface | 2376 | 2657 | 20693 | 2215.16+-1544.44 | 1050.37+-510.51 |
| I2 | 6800STDY12499506 | PD45816 | Kidney - Tum. Interface | 2376 | 1777 | 21399 | 5384.21+-2836.88 | 2242.85+-873.76 |
| I5 | 6800STDY12499508 | PD47465 | Kidney - Tum. Interface | 2376 | 2609 | 21840 | 3565.95+-2075.09 | 1520.86+-650.56 |
| I3 | 6800STDY12499406 | PD43824 | Kidney - Tum. Interface | 2376 | 3153 | 22149 | 2479.87+-2041.36 | 1191.62+-657.54 |
| I1 | 6800STDY12499411 | PD47171 | Kidney - Tum. Interface | 2376 | 2048 | 23116 | 8371.10+-5489.16 | 2993.72+-1393.52 |

#### 5.1.2 CRC dataset

We included in this study a primary tumor biopsy (*PT*) and two liver metastases (*LM4* and *LM7*). Biopsies have been collected in the Department of Oncology University of Leuven from a single patient (*SC087*). For LM4, we generated a technical replicate (*LM4r*). Slices from *PT* and *LM7* were smaller than LM4, so it was possible to put two consecutive slices on the same stereo-seq chip.

##### Donors and Sample collection

Biopsies from patients diagnosed with Colorectal Cancer (CRC) and liver metastasis were received fresh after written informed consent, according to the Declaration of Helsinki. Approval by the medical ethics commission of the KU Leuven University (S50887) was obtained, in accordance whith the local ethical guidelines. Samples were collected from a 72 years old male patient. (*SC087*). Before surgery, patient received a combination of immunotherapy (i.e., Cetuximab) and chemotherapy (i.e., Levofolinezuur, Oxaliplatine plus Fluorouracil) treatment.

##### Histological Analysis

To assess the morphological features and tissue structure and preservation, we performed Hematoxylin and Eosin (H&E) staining of each tissue block. The fresh frozen samples were sectioned at a thickness of 10*µm* and fixed with methanol. Afterward, sections were incubated with 500*µl* of isopropanol for 1 minute. Slides were air-dried for 5-10 minutes, followed by incubation with hematoxylin. Afterward, the slides were washed by immersion in water and air-dried. Subsequently, 1 ml of bluing buffer was applied and washed. Eosin mix was then added to stain the cytoplasmic components and extracellular matrix. After washing, the stained sections were imaged in a microscope using a BF-Epi channel, 4X and 10X objective lenses, with stitching function.

##### Stereo-seq Tissue Optimization

Tissue optimization was performed using the Stereo-seq Permeabilization kit (Cat. No. 111KP118) and Stereo-seq chip set P (Cat. No. 110CP118) according to the manufacturer’s protocol (Stereo-seq permeabilization set user manual, Ver A1) to define the optimal permeabilization time for the tissue samples we specifically wanted to analyze. Briefly, 10*µm* tissue sections were prepared from the primary tumor and liver metastasis cryo-blocks and placed on 2 permeabilization slides containing 4 chips each. The tissue layer was thawed to attach it to the surface of the chips. After drying the tissue at 37^*o*^*C*, the slides were then dipped into pre-chilled 100% methanol at − 20^*o*^*C* and incubated for 30^*o*^*C* min to fix the tissue. Post fixation, the tissue permeabilization test was performed on these chips by permeabilizing the tissue at 4 different time points (6, 12, 18, and 24 minutes). Afterward, reverse transcription was carried out at 42^*o*^*C* for 1 h in dark, followed by tissue removal at 55^*o*^*C* for 1 h. Fluorescence imaging was performed in the TRITC channel with 10X objective, following the imaging guidelines provided by the manufacturer (Guidebook for Image QC & microscope assessment and imaging, Ver A5). The optimal permeabilization time was assessed based on the strongest fluorescence signal with the lowest signal diffusion (crispness of the RNA footprint). We found the optimal permeabilization time for both the primary colon tumor and the liver metastasis to be between 12 and 18 min and, thus, 15 min was used for the spatial transcriptomics assay.

##### Stereo-Seq Spatial Transcriptomics Assay

The spatial transcriptomics analysis was performed using the Stereo-seq Transcriptomics kit (Cat. No. 111ST114) according to the manufacturer’s protocol (Stereo-seq Transcriptomics set user manual, Ver A2). Briefly, cryosectioning, tissue mounting, and fixation were performed exactly as previously described in the protocol for the tissue optimization. Next, each fixed tissue section was stained with Qbit ssDNA reagent. Fluorescence imaging of the single-stranded DNA staining was performed in the FITC channel with a 10X objective, following the imaging guidelines provided by the manufacturer (Guidebook for Image QC & microscope assessment and imaging, Ver A5). Prior to permeabilization, the ssDNA-stained image was also subjected to QC analysis using the imageQC software as recommended by the manufacturer. Tissues were then permeabilized for 15 minutes, as estimated in the tissue permeabilization test.

After washing the chip, reverse transcription was performed at 42 for 3 h. The tissue was then digested and removed from the chip and cDNA was released, collected, and purified following the manufacturer’s recommendation. After quality assessment using a bioanalyzer (Agilent), sequencing library preparation was performed using transposase assisted tagmentation reaction. Indexed PCR and library purification were performed to prepare the final sequencing library as per manufacturer’s recommendations. Final Stereo-seq libraries were sequenced on MGI/BGI sequencing platforms (DNBSEQTM T7) at the MGI Latvia sequencing facility.

##### Stereo-Seq Sequencing Processing

We processed each FASTQ file obtained from each sample using the SAW pipeline v.7.1.1 [44]. The pipeline consists of two main steps, which were performed for each sample as follows: 1) *Alignment and Counting* : We used Homo sapiens.GRCh38.dna.primary assembly.fa, Ensembl release 111, as the reference genome and the corresponding GTF annotation file (Homo sapiens.GRCh38.111.gtf). The sample-related chip mask file, which includes the spatial coordinates, was part of the transcriptomic kit provided by STOmics. We reported the sample IDs in Table 1. 2) *Image Registration*: The fluorescence image taken during the *Stereo-Seq Spatial Transcriptomics Assay* was aligned to the count matrix to define the tissue area, filtering out DNA nano-balls that were not under the tissue from the count matrix.

The pipeline outputs the registered image and a table (.gem/.gef) that includes, for each unique combination of gene ID and spatial coordinates (x, y), the number of molecular identifier (MID) counts. This table was further processed with Stereopy v1.3.0 [45] to define the final count matrix by aggregating spatial coordinates into spots of 50 × 50 DNA nano-balls. Therefore, the dimension of each spot is 25 × 25*µm*^2^. The obtained count matrix (spot×gene) is stored in a Scanpy object for downstream analyses.

### 5.2 ST read counts preprocessing

We applied the same preprocessing steps for both Visium and Stereo-seq datasets using Scanpy v1.9.6 [46]. In particular, the expression profile in each spot was normalized by library size (*pp*.*normalize total(adata=adata, target sum=1e4)*) and log-transformed (*pp*.*log1p(adata)*) as usually done in single-cell data analysis [47].

Subsequently, we ran Principal Component Analysis (*tl*.*pca(adata, svd solver=‘arpack’)*) and selected the first 30 PCs as input for k-nearest-neighbor graph (kNN) construction (*pp*.*neighbors(adata, n neighbors=10, n pcs=30)*).

Next, we clustered kNN graph nodes using the Leiden approach (*tl*.*leiden(adata)*), selecting a resolution specific to each dataset. We explored the data quality by plotting the distribution of detected genes and total counts in each cluster via violin plots. If needed, we removed clusters with particularly low numbers of detected genes and counts. The resulting datasets have the dimensions reported in table 1.

Finally, to reduce data sparsity, we applied MAGIC imputation [48] with default parameters, before processing the datasets with spFBA.

### 5.3 Metabolic model

spFBA theoretically accepts any metabolic network model. However, we were more confident using the manually curated, recently published ENGRO2 core network model [14] of the human central carbon and essential amino acids metabolism. ENGRO2 includes 484 reactions, 403 metabolites, and 497 genes, and represents a follow-up of the core model of human central metabolism ENGRO1 [35]. For this model, 337 reactions are associated with a GPR rule. The biomass pseudo-reaction corresponds with the biomass reaction of the Recon3D model, in terms of the set of metabolites considered and corresponding stoichiometric coefficients. We simulated a growth medium condition in which exogenous metabolites are abundantly available, or open medium. Exogenous metabolites in the ENGRO2 network include glucose, lactate, oxygen, water, hydrogen, folic acid, and all essential and non-essential amino acids.

### 5.4 spFBA methodology

#### 5.4.1 Reaction Activity Scores

We computed a Reaction Activity Score (RAS) for each reaction and spot by substituting the mRNA abundances into the corresponding Gene-Protein-Reaction (GPR) rule, as done in [49]. To solve the logical expressions, the minimum transcript value was taken when multiple genes are joined by an AND operator, and the sum of their values was taken when multiple genes are joined by an OR operator. In the case of GPRs combining both operators AND and OR, we respected their standard precedence.

Once a RAS was computed for each reaction in the metabolic model, we obtained a table of dimensions Spots x Reactions, containing the RAS values.

#### 5.4.2 Flux Variability Analysis

To determine the extreme points of the feasible space, that is, the range of possible fluxes that satisfy the mass balance and the medium constraints, we used Flux variability analysis (FVA)[50, 51, 52]. FVA is a constraint-based modeling technique aimed at determining the maximal and minimal possible flux through any reaction of the model. FVA solves the following two linear programming optimization problems (one for minimization and one for maximization) for each flux *v*_*r*_ of interest, with *r* = 1,. .., *R*:

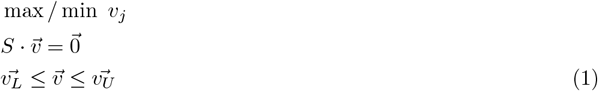

where *S* is the stoichiometric matrix provided by a metabolic network model, 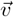 is the vector representing the flux of each reaction, 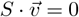 represents the steady state assumption, and the vectors 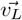 and 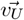 representing the flux boundaries used to mimic as closely as possible the biological process in the analysis and the availability of nutrients in the medium define the feasible region of steady-state flux distributions.

We assumed a rich open medium where all exogenous metabolite are made available in an unlimited quantity, which is 1000 in the model.

#### 5.4.3 Spot-relative flux constraints

After the RAS and FVA computation, specific constraints on internal fluxes of the network are built following an approach adapted from [14, 15]. For each reaction *j* = 1,. .., *R* and spot *s* = 1,. .., *S*, an upper bound 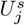 and a lower bound 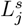 to the flux capacity are defined, based on the following formulas:

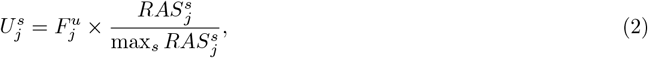

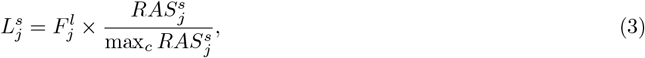

where 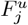 and 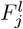 represent the maximum and the minimum flux that reaction *j* might carry, obtained by FVA, and 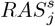 is the RAS value for spot *s* and reaction *j*. These constraints are used to map the transcriptomics data into spot lattice, with a one-to-one correspondence between the single spot transcriptomics profile and the corresponding network of each spot. Therefore, each sub-network has specific constraints derived from the transcriptomics and defined in Eq. 2 and 3.

#### 5.4.4 Parsimonious FBA

Given the new set of spot-relative constraints for each spot, we used Eq. 1 to determine the flux distribution that optimizes metabolic growth (i.e., the flux of the biomass synthesis pseudo-reaction) for each spot. Because the optimization problem might have alternative solutions, to select a single flux distribution we employed parsimonious FBA (pFBA). pFBA operates in two optimization steps. First, it determines the maximal value of the biomass. Then, it minimizes the total sum of reaction fluxes while maintaining the optimal value of the primary objective [53].

#### 5.4.5 Corner-based sampling

To analyze each spot-relative metabolic network assuming an objective function, spFBA relies on flux sampling of the feasible region to generate a sequence of feasible solutions that satisfy the network constraints. Flux sampling provides information not only on the range of feasible flux solutions, like FVA, but also on their probability.

In this study, we employed an algorithm, called Corner-based Strategy (CBS) [54, 55], to sample the vertices of the feasible region by using different weighted objective functions each time as the objective function. A predefined number of samples is determined. At each iteration, a new objective function is set by randomly assigning weights in the range [−1, 1] and deciding whether the objective function will be maximized or minimized. To account for the different scales of the various reactions in the model, the weights are divided by the FVA maximum value for each specific reaction. More details of the implementation are reported in[55]. Such an algorithm captures flux distributions qualitatively and quantitatively more heterogeneous compared to classical Hit-and-Run strategies. In this work, we sampled 10000 points for each spot.

#### 5.4.6 Flux Enrichment Score (FES)

For each spot, we calculated a single Flux Enrichment Score (FES) using the following approach. For the pFBA method, we obtained the unique flux distribution by solving the corresponding optimization problem. We then normalized each reaction value by taking the maximum of the FVA if the flux value is positive, and the minimum of the FVA otherwise. This normalization allows us to assign a score between −1 and 1 for each reaction. In the case of the CBS method, we computed the centroid from 1000 samples, which yields a feasible flux distribution based on the average of all sampled fluxes. We then applied the same normalization process as in the pFBA method. For each FES we also computed its 99% confidence interval.

#### 5.4.7 Spots clustering

For each layer of features, namely counts, RAS, and the 2 different types of fluxes (pFBA ad CBS), we first applied Principal Component Analysis (PCA) to reduce the dimensionality of the datasets. The neighborhood graph was computed on the PCA space to create a distance matrix, encoding the connectivity between spots based on Euclidean distance [56]. The Leiden community detection algorithm was then performed to identify the optimal communities representing clusters of well-connected spots within the neighborhood graph [57].

To produce comparable clusters, we optimized the clustering parameters by maximizing the silhouette score [58], by testing different parameter combinations with a grid-search strategy. The parameters optimized included the number of principal components, the number of neighbors when computing the neighborhood graph, and the resolution in the Leiden algorithm.

## 6 Resource availability

### 6.1 Lead Contact

Requests for further information and resources should be directed to and will be fulfilled by the lead contact, Chiara Damiani.

### 6.2 Materials availability

This study did not generate new unique reagents.

### 6.3 Data and code availability

The Python implementation of the spFBA computational pipeline, the code for generating the main figures, and the raw and preprocessed read counts datasets including both Kidney (10x Visium), Colon, and Liver (Stereo-seq) samples, are available in the following Zenodo repository: *DOI: 10*.*5281/zenodo*.*13988865*. The repository also includes the main output files. In particular, the FESs for all the samples.

The repository will be made public upon publication. Private link for reviewers: https://zenodo.org/records/13988866?preview=1&token=eyJhbGciOiJIUzUxMiJ9.eyJpZCI6IjU0MzhjNTE4LTlmMTItNDg5ZS1iZmQ1LTBkNTI5NjgyOGJhNiIsImRhdGEiOnt9LCJyYW5kb20iOiJkZDhjNmJmMjUzNmVhNjA5ZmMzMWU1ZDEwNTVkYzNiMCJ9.5ZZZAzIwit3HoWLt-EUJ6ShFx3XW8M9BIrlV2ZhrFrhglmChZCpKO90pHIa2anB9gJjfPXZYqMJR8Ojc-U8w.

## 8 Acknowledgments

We sincerely thank Zhigang Lu and his team for preparing and sequencing the CRC samples. We are also deeply grateful to the patients who generously consented to provide tissue samples for this study, making this work possible. We would also like to thank Marco Vanoni for his valuable input during insightful discussions that contributed to this work.

## 9 Author contributions

Conceptualization, D.M., A.G., H.H, and C.D.; Methodology, G.M., F.L., B.G.G., and C.D.; Software, D.M., G.M., F.L., and B.G.G.; Formal Analysis, D.M., G.M., F.L., B.G.G., and C.D.; Investigation, D.M., G.M., F.L., B.G.G., and I.R.; Resources, B.V., K.Y., and S.T.; Data Curation, D.M., F.L., I.R., and K.Y.; Writing – Original Draft, D.M., G.M., B.G.G., and C.D.; Writing – Review & Editing, D.M., S.T., A.G., H.H., A.P.R., and C.D.; Visualization, D.M., G.M., I.R., and A.G.; Supervision, S.T., H.H., A.P.R., and C.D.; Project Administration, S.T., H.H., and C.D.; Funding Acquisition, D.M., S.T., H.H., and C.D.;

