## Supplementary material for "Spatial Flux Balance Analysis reveals region-specific cancer metabolic rewiring and metastatic mimicking": Supplemetary information

### Supplementary Materials

#### Supplementary Figures

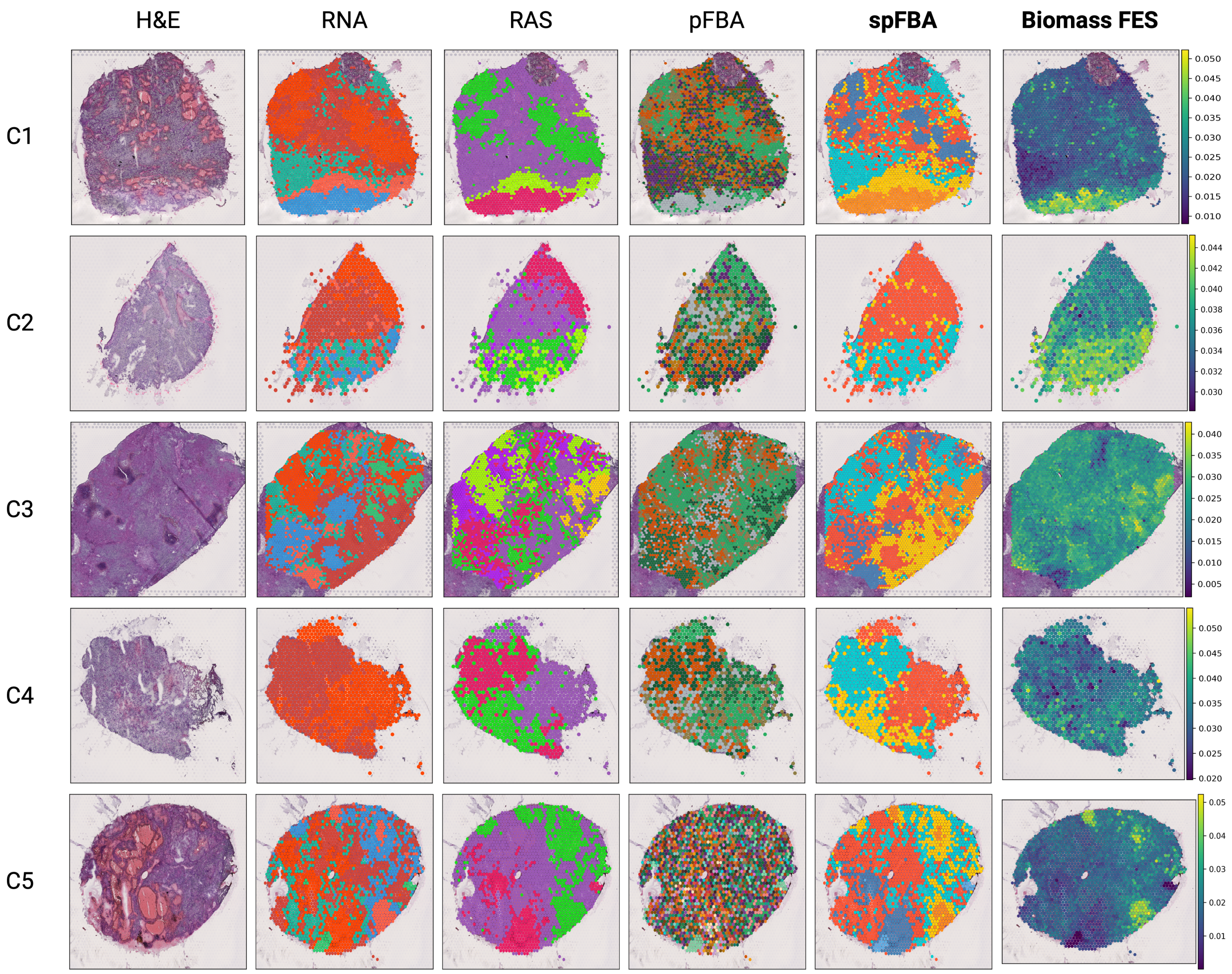


Figure S1: ccRCC tumor core samples. From left to right: H&E staining; clustering results obtained with different data layers: RNA (processed reads counts), RAS, FESs obtained with pFBA and spFBA - colors were arbitrarily assigned to clusters, thus same colored clusters in different plots are not directly comparable; Biomass FES obtained with spFBA

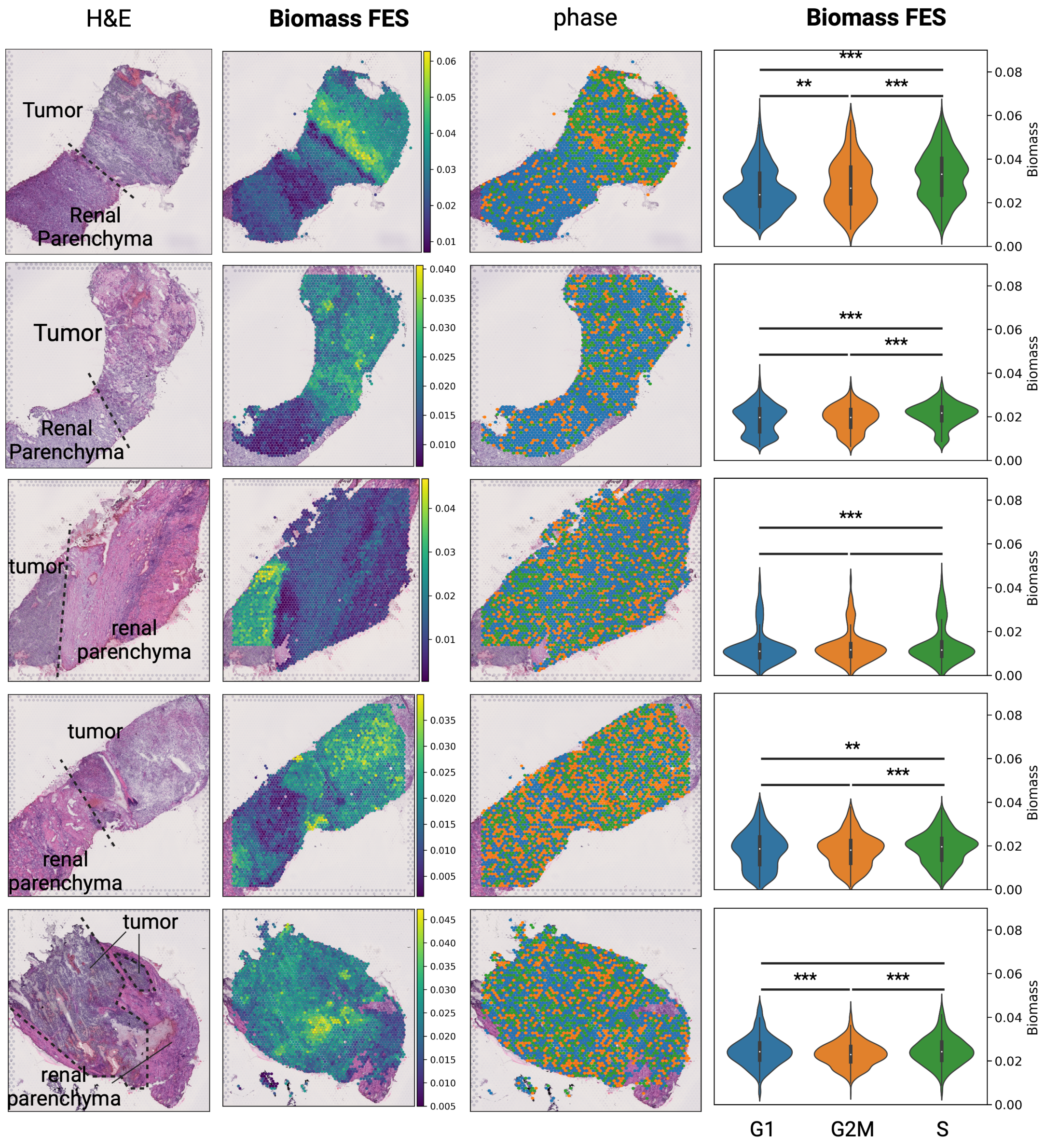


Figure S2: From left to right: H&E staining highlighting the division between tumor and renal parenchyma; Spatial distribution of Biomass FES; Spatial distribution of the most likely cell-cycle phase and frequency distribution of the biomass FES for the spots at each phase [2]. It can be observed that the biomass FESs tend to be higher (Mann-Whitney $p < 0.05$) in spots classified as being in the S phase compared to those in the G1 phase for most of the samples [Figure 2]. The S phase of the cycle is characterized by DNA replication, indicating active cell proliferation. We must however remark that a cell classified as being in the G1 phase does not necessarily exclude it as proliferative. Moreover, spots do not have single-cell resolution and it is reasonable to assume that the cells within a single spot are not synchronized in their cell cycles. Therefore, the final cell-cycle score for each spot is a combination of the individual cell's scores within that spot. Nevertheless, spots classified as S phase are indicative of the presence of proliferative cells.

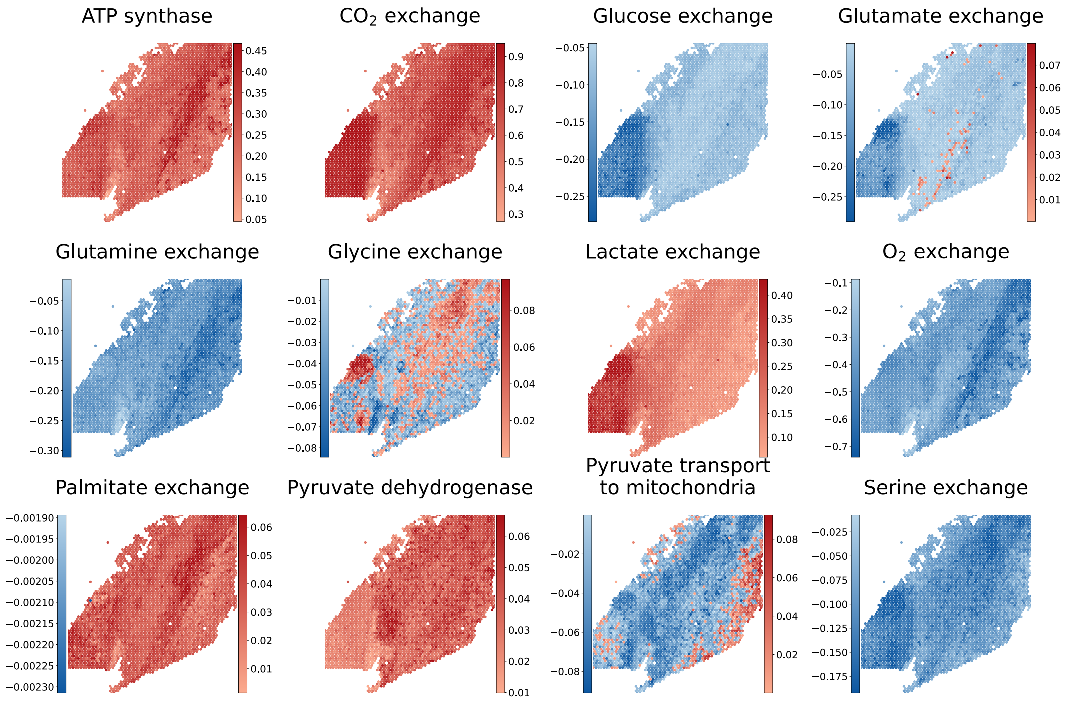


Figure S3: The FES of a set of reactions of interest (in alphabetic order) is visualized for ccRCC sample I3. In the case of exchange reactions, negative values correspond to consumption of the metabolite, positive values to production.

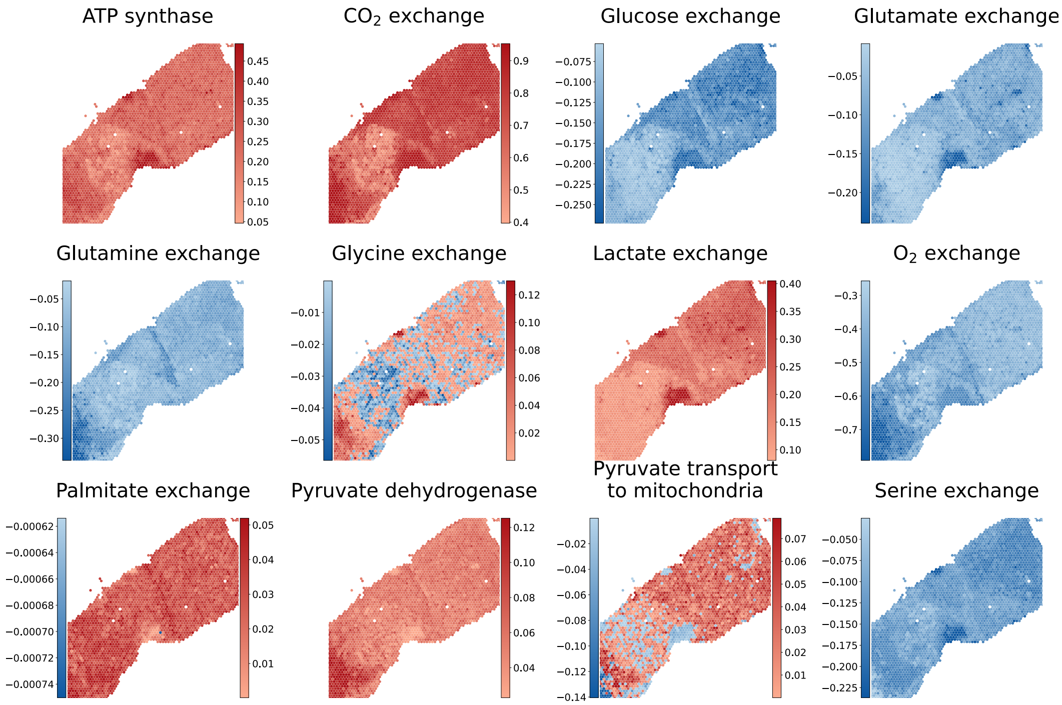


Figure S4: The FES of a set of reactions of interest (in alphabetic order) is visualized for ccRCC sample I4. In the case of exchange reactions, negative values correspond to consumption of the metabolite, positive values to production.

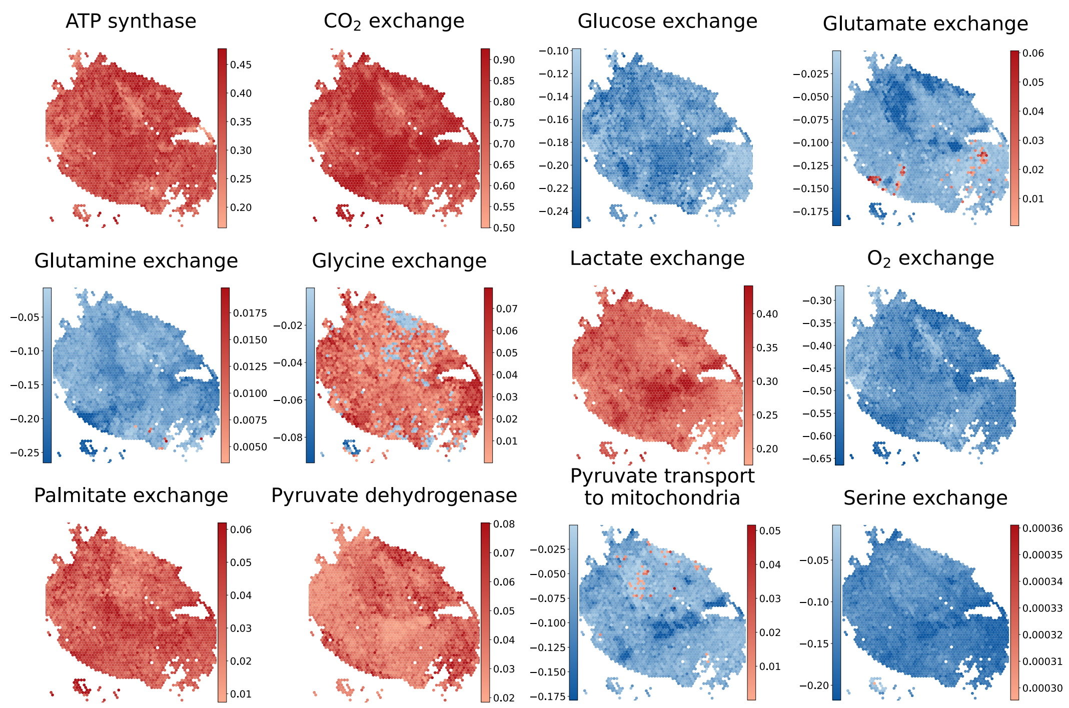


Figure S5: The FES of a set of reactions of interest (in alphabetic order) is visualized for ccRCC sample I5. In the case of exchange reactions, negative values correspond to consumption of the metabolite, positive values to production.

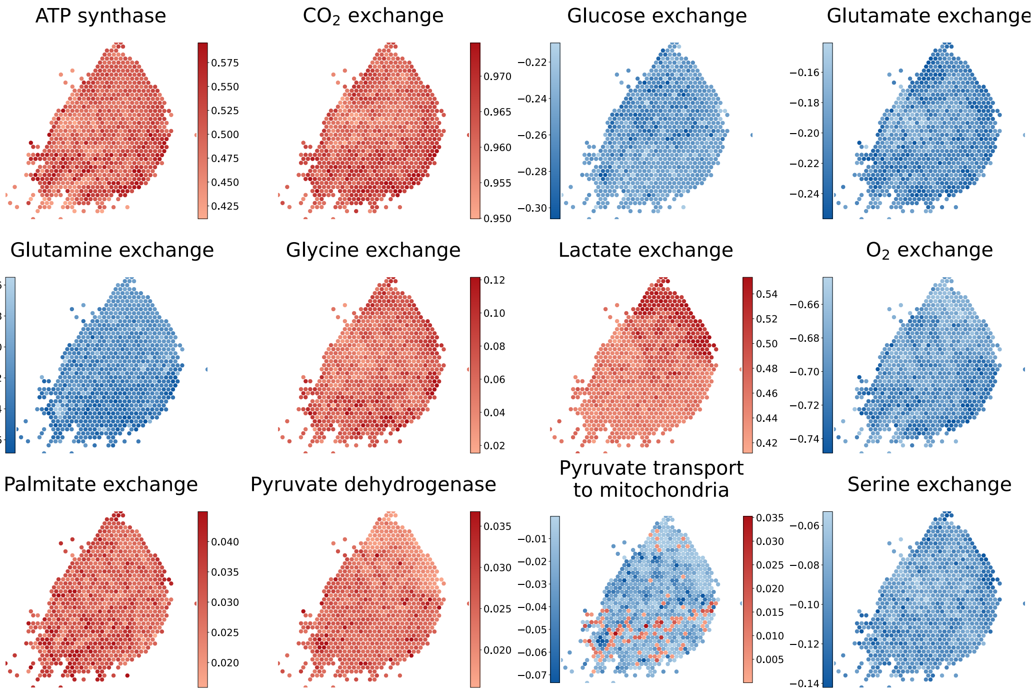


Figure S6: The FES of a set of reactions of interest (in alphabetic order) is visualized for ccRCC sample C2. In the case of exchange reactions, negative values correspond to consumption of the metabolite, positive values to production.

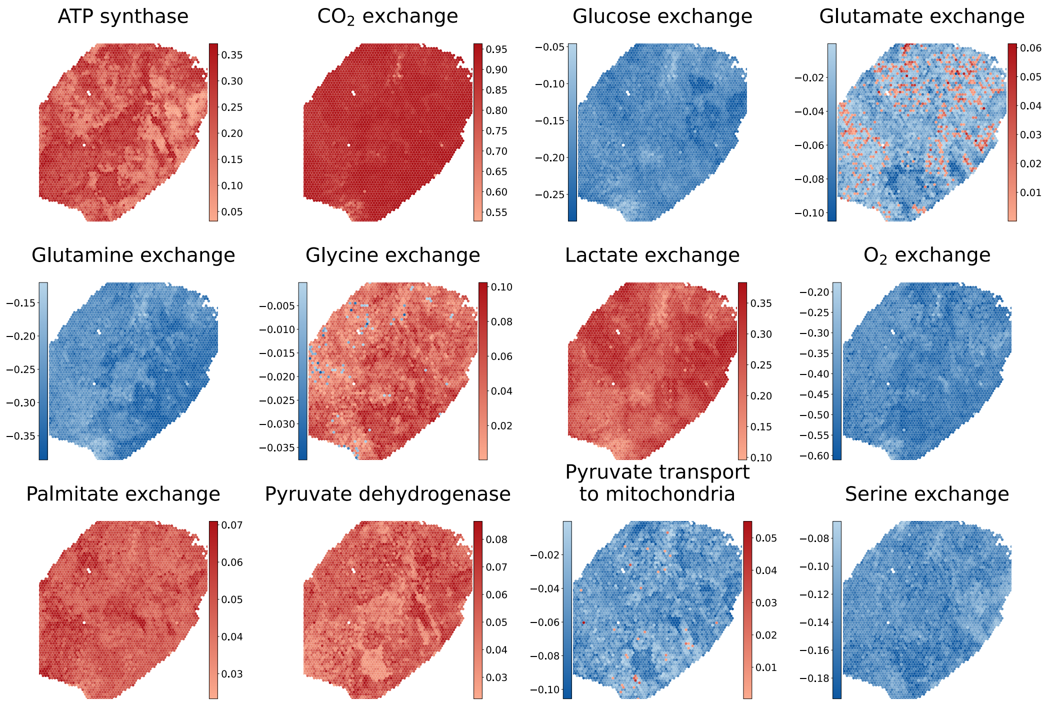


Figure S7: The FES of a set of reactions of interest (in alphabetic order) is visualized for ccRCC sample C3. In the case of exchange reactions, negative values correspond to consumption of the metabolite, positive values to production.

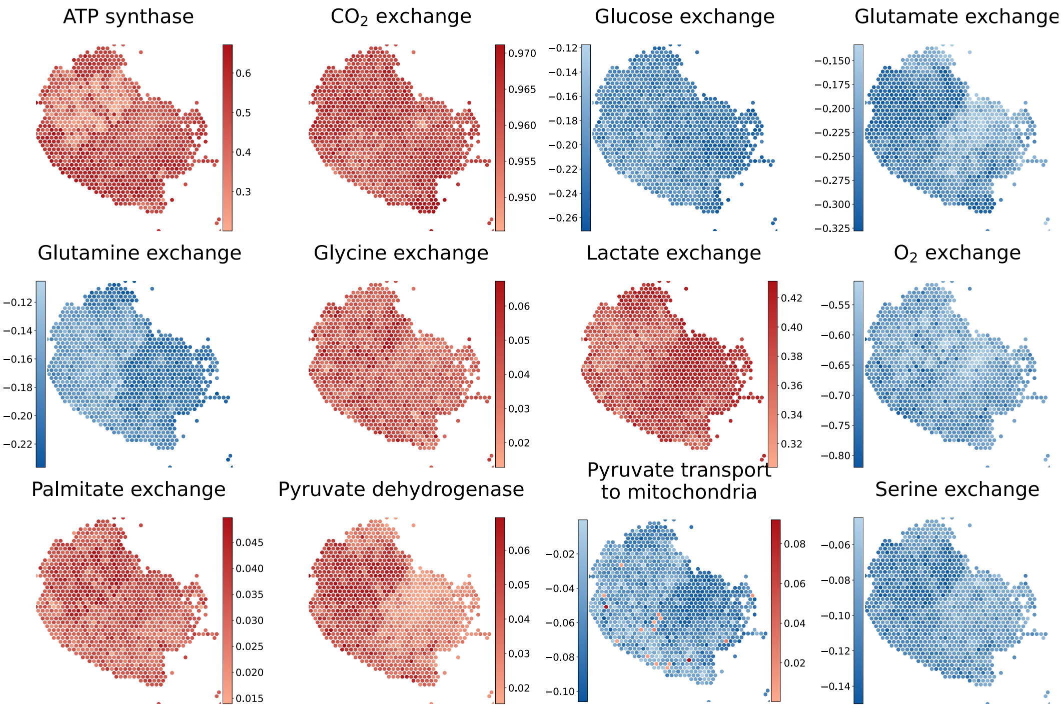


Figure S8: The FES of a set of reactions of interest (in alphabetic order) is visualized for ccRCC sample C4. In the case of exchange reactions, negative values correspond to consumption of the metabolite, positive values to production.

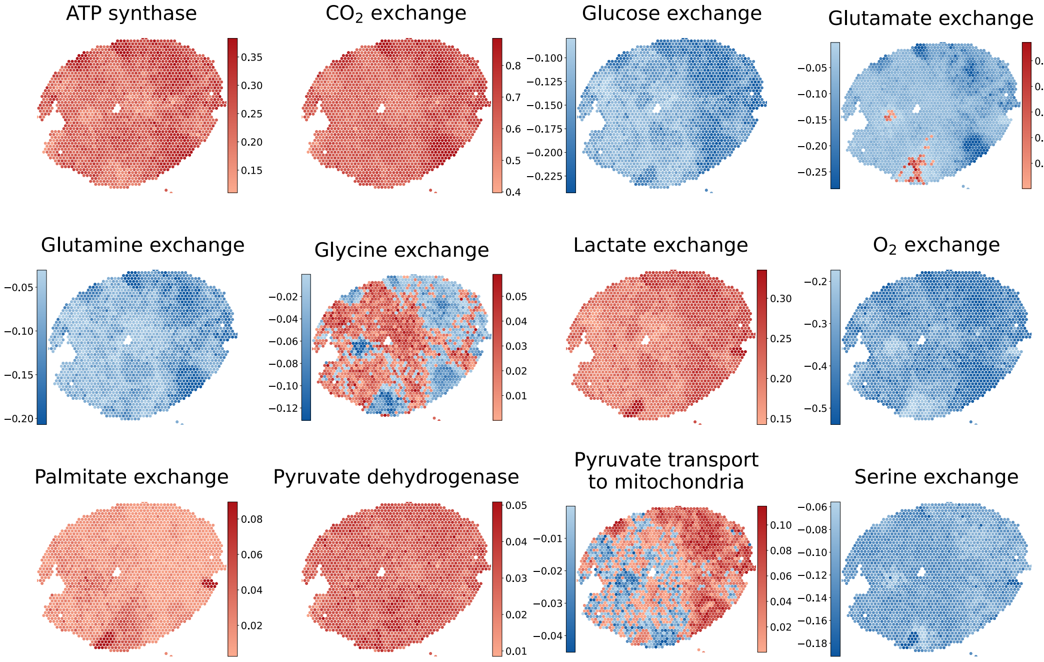


Figure S9: The FES of a set of reactions of interest (in alphabetic order) is visualized for ccRCC sample C5. In the case of exchange reactions, negative values correspond to consumption of the metabolite, positive values to production.

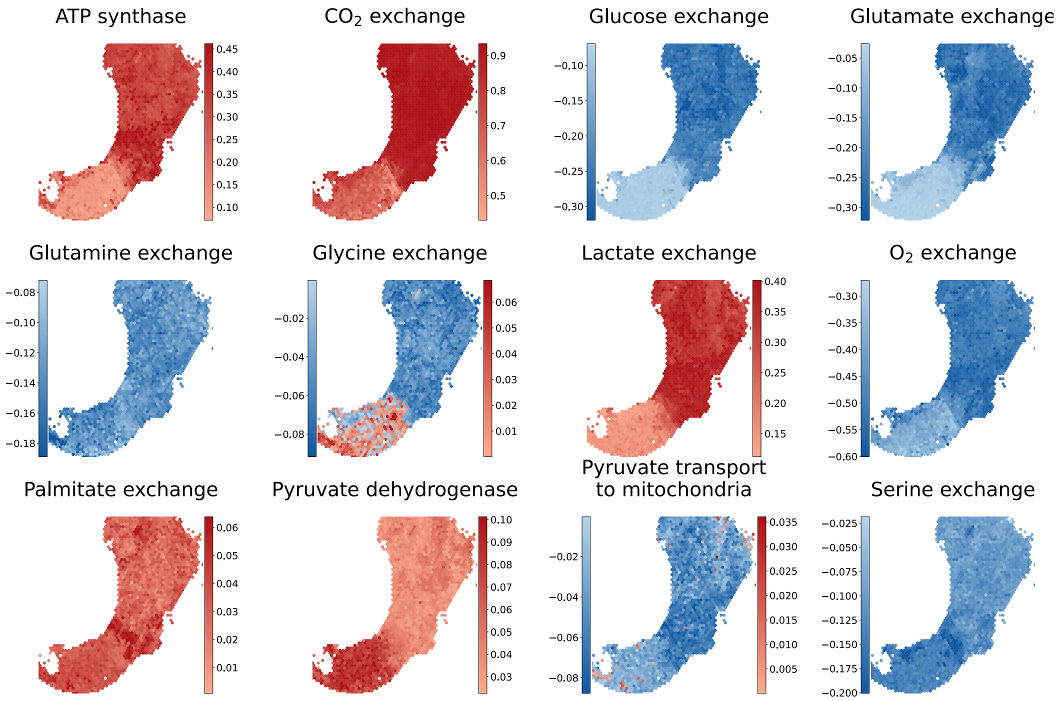


Figure S10: Lower bound of the 99% confidence interval for the FES of a set of reactions of interest (in alphabetic order) is visualized for ccRCC sample I2. In the case of exchange reactions, negative values correspond to consumption of the metabolite, positive values to production.

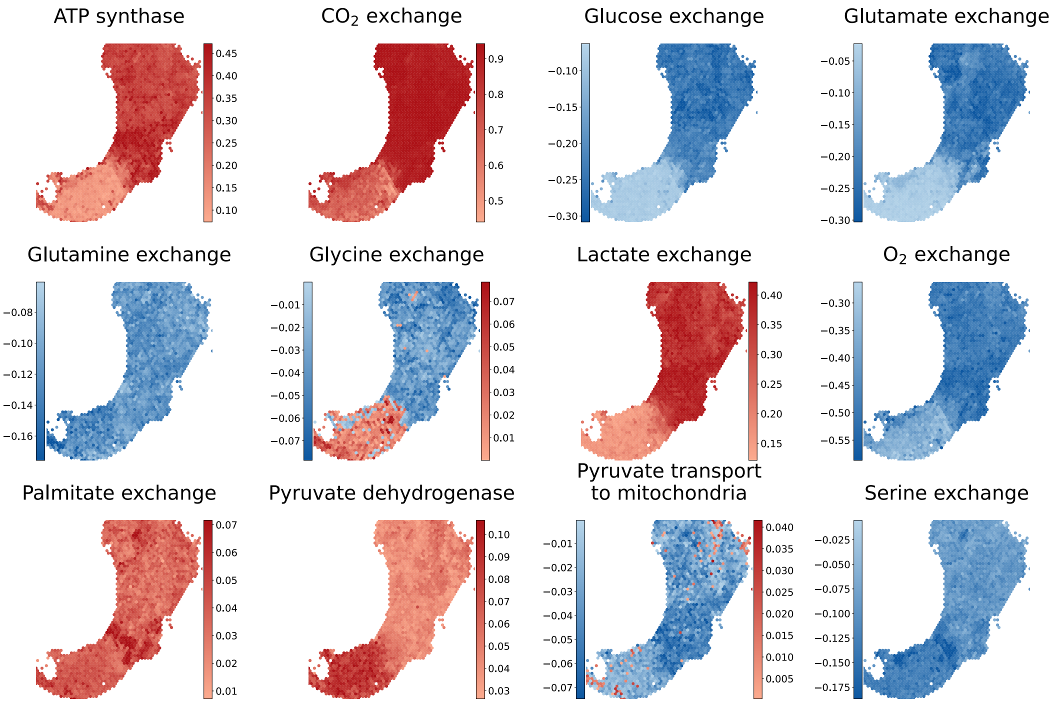


Figure S11: Upper bound of the 99% confidence interval for the FES of a set of reactions of interest (in alphabetic order) is visualized for ccRCC sample I2. In the case of exchange reactions, negative values correspond to consumption of the metabolite, positive values to production.

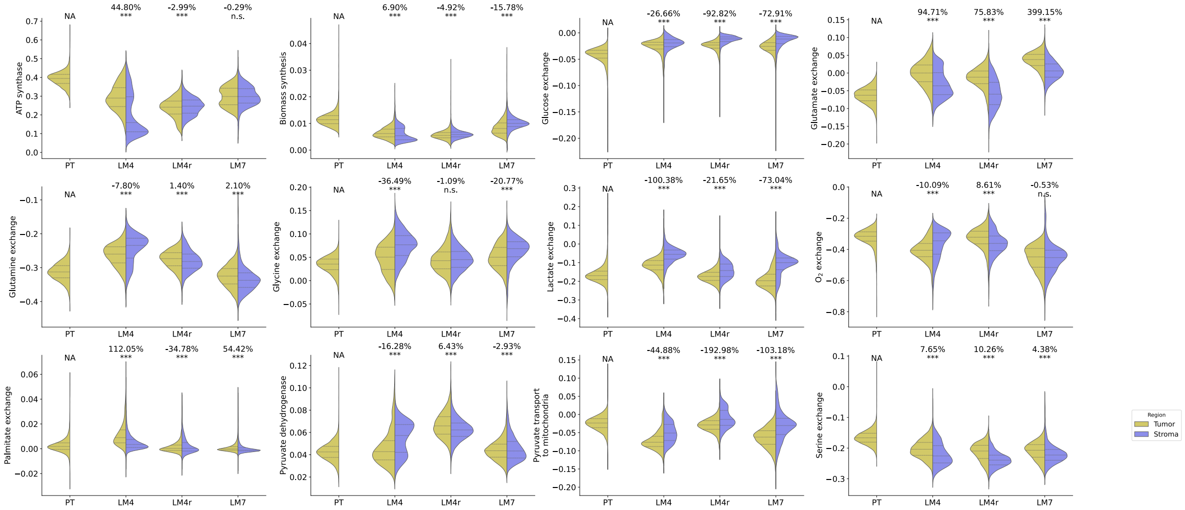


Figure S12: Statistical comparison of the FESs between spots annotated as stroma and those annotated as tumor (based on annotations in Fig. S15) for the selected reactions, listed in alphabetical order. The p-values were calculated using t-tests

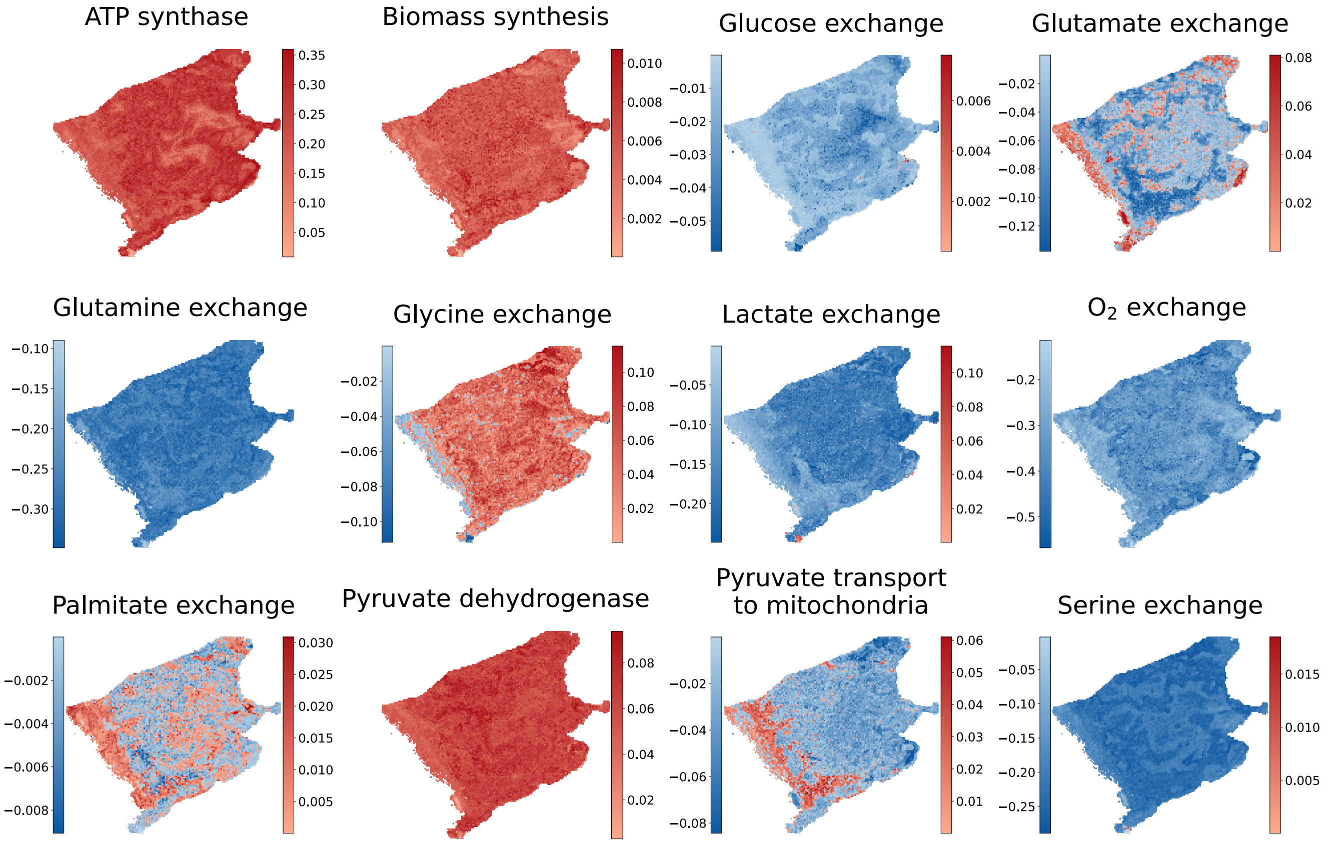


Figure S13: FES of a set of reactions of interest (in alphabetic order) for CRC LM4r. In the case of exchange reactions, negative values correspond to consumption of the metabolite, positive values to production.

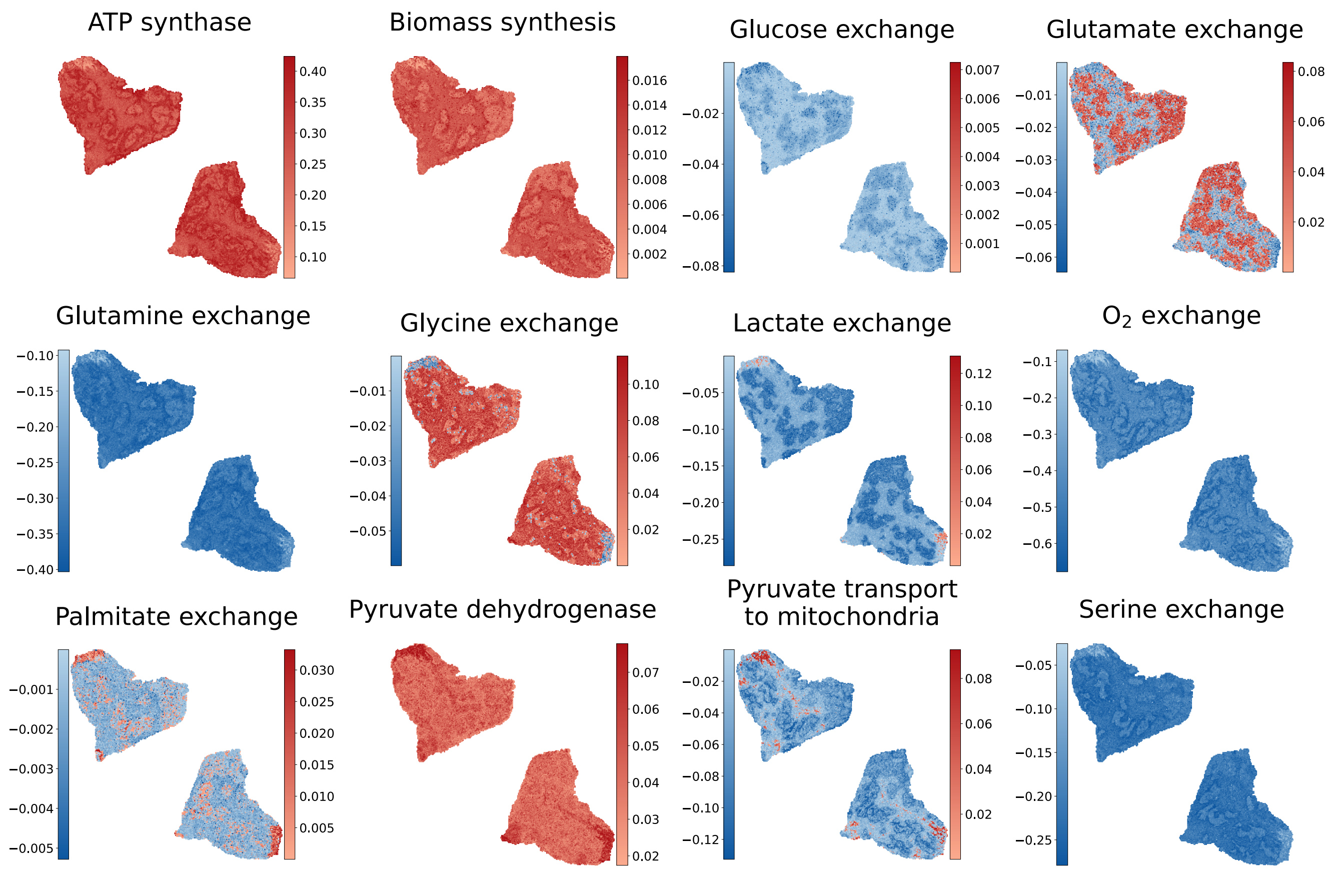


Figure S14: FES of a set of reactions of interest (in alphabetic order) for CRC LM7. In the case of exchange reactions, negative values correspond to consumption of the metabolite, positive values to production.

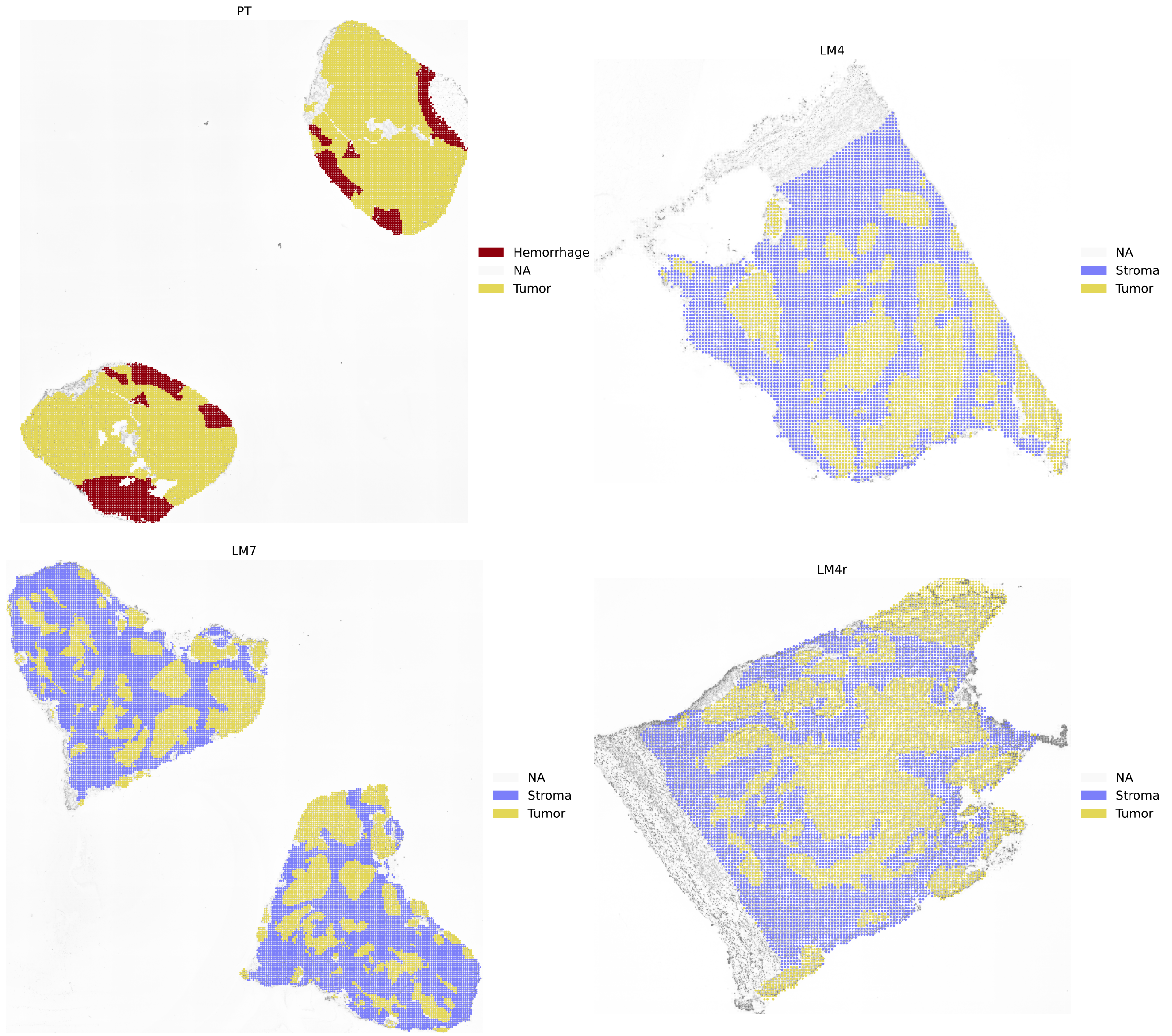


Figure S15: H&E stained images of the subsequent slice of each colorectal cancer sample. Three main regions were annotated using QuPath 0.5.1. Images were manually rotated to match the sequenced slice positions.

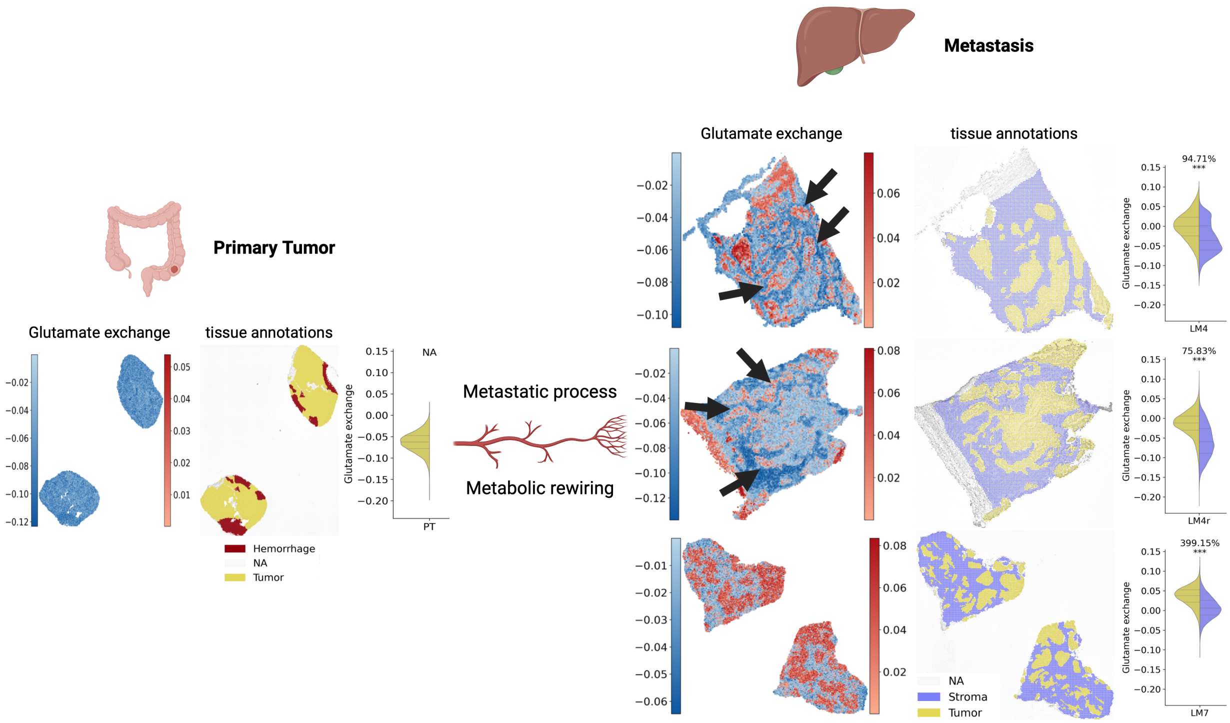


Figure S16: On the left, the violin plot represents glutamate exchange in the primary tumor (PT) region, where stromal tissue is absent, showing uniform glutamate consumption across the tumor without statistical testing. On the right, three metastatic liver samples (LM7, LM4, LM4r) are shown. For each metastasis, spatial maps of glutamate exchange (center) reveal heterogeneity within the tumor region. Black arrows indicate areas at the tumor-stroma interface where glutamate production is detected, in contrast to core regions that predominantly consume glutamate, albeit at a lower rate than stromal cells. Adjacent to the spatial maps, hematoxylin and eosin (H&E)-based tissue annotations highlight tumor (yellow) and stroma (blue). The violin plots on the far right compare stromal and tumor regions for each metastatic sample, with statistical tests indicating significant differences in glutamate exchange between the two compartments

#### Supplementary Tables

| | **spFBA** | **pfBA** | **RAS** | **RNA** |
|---|---|---|---|---|
| **spFBA** | - | - | - | - |
| **pfBA** | 0.28 | - | - | - |
| **RAS** | 0.58 | 0.29 | - | - |
| **RNA** | **0.55 ***** | 0.30 | 0.66 | - |

Table S1: Mean V-Measures across data layers. To compare the clustering analysis results across layers, we employed the V-measure. V-measure is an entropy-based cluster quality evaluation score given the class labels [59]. After computing the V-measure score, by comparing the clustering results from each layer against each other as reference class labels, a table of all the V-measures was calculated. The mean V-measure across all samples was then computed. The results indicate that spFBA display a significantly higher V-measure than pFBA, demonstrating that spFBA more consistently reflects the tissue architecture represented by the clusters at the normalized counts level (RNA). A Mann-Whitney U test comparing the means showed that spFBA had a significantly higher mean V-measure compared to pFBA ($p < 0.05$).
